# Evolutionary Divergence in the Conformational Landscapes of Tyrosine vs Serine/Threonine Kinases

**DOI:** 10.1101/2022.08.29.505757

**Authors:** Joan Gizzio, Abhishek Thakur, Allan Haldane, Ronald M. Levy

## Abstract

Inactive conformations of protein kinase catalytic domains where the DFG motif has a “DFG-out” orientation and the activation loop is folded present a druggable binding pocket that is targeted by FDA-approved “type-II inhibitors” in the treatment of cancers. Tyrosine Kinases (TKs) typically show strong binding affinity with a wide spectrum of type-II inhibitors while Serine/Threonine Kinases (STKs) usually bind more weakly which we suggest here is due to differences in the folded to extended conformational equilibrium of the activation loop between TKs vs. STKs. To investigate this, we use sequence covariation analysis with a Potts Hamiltonian statistical energy model to guide absolute binding free-energy molecular dynamics simulations of 74 protein-ligand complexes. Using the calculated binding free energies together with experimental values, we estimated free-energy costs for the large-scale (∼17-20Å) conformational change of the activation loop by an indirect approach, circumventing the very challenging problem of simulating the conformational change directly. We also used the Potts statistical potential to thread large sequence ensembles over active and inactive kinase states. The structure-based and sequence-based analyses are consistent; together they suggest TKs evolved to have free-energy penalties for the classical “folded activation loop” DFG-out conformation relative to the active conformation that is, on average, 4-6 kcal/mol smaller than the corresponding values for STKs. Potts statistical energy analysis suggests a molecular basis for this observation, wherein the activation loops of TKs are more weakly “anchored” against the catalytic loop motif in the active conformation, and form more stable substrate-mimicking interactions in the inactive conformation. These results provide insights into the molecular basis for the divergent functional properties of TKs and STKs, and pharmacological implications for the target selectivity of type-II inhibitors.

## Introduction

The human genome contains approximately 500 eukaryotic protein kinases which coordinate signaling networks in cells by catalyzing the transfer of a phosphate group from ATP to serine, threonine, or tyrosine residues^1,2^. The GO (Gene Ontology) database identifies 351 (∼70%) of these enzymes as Serine/Threonine Kinases (STKs), and 90 (∼18%) as Tyrosine Kinases (TKs). STKs are an ancient class of protein kinases that predate the divergence of the three domains of life (bacteria, archaea, eukaryote)^3^, whereas TKs are a more recent evolutionary innovation, having diverged from STKs about 600 million years ago during early metazoan evolution^4,5^. Kinases are important therapeutic targets in a large number of human pathologies and cancers. Both TKs and STKs share a striking degree of structural similarity in their catalytic domains, owing to evolutionary selective pressures that preserve their catalytic function; in particular, the location and structure of the ATP binding site is highly conserved which raises significant challenges for the design of small-molecule ATP-competitive inhibitors that are both potent for their intended target(s) and have low off-target activity for unintended kinase targets. The latter is referred to as the “selectivity” of an inhibitor, a property which is difficult to predict and control but is nonetheless very important for developing drugs with minimal harmful side effects.

A particular class of ATP-competitive kinase inhibitors which were proposed to have a high potential for selectivity are called “type-II inhibitors” which only bind when the kinase adopts an inactive “DFG-out” conformation. “DFG” is a conserved catalytic motif located at the N-terminus of a ∼20 residue activation loop that is highly flexible and controls the activation state of the kinase and the structure of the substrate binding surface. In both TKs and STKs, this “activation loop” undergoes a large-scale (∼17Å) conformational change when the DFG motif flips from the active “DFG-in” conformation where the activation loop is “extended”, to the classical DFG-out conformation where the activation loop is “folded”. The flip of the DFG motif from “in” to “out” opens a back pocket which is connected to the conserved ATP binding site through a “gatekeeper” residue. Type-II inhibitors typically have a long chemical fragment that allows them to bind across the gatekeeper and form interactions with residues in the “back pocket” which is formed due to the DFG flip. In contrast, type-I inhibitors (the majority of kinase drugs) occupy the ATP pocket but not the back pocket and can bind to either DFG-in or DFG-out. For these reasons, it has been proposed that type-II inhibition holds greater potential for the design of highly selective drugs^6–8^; it has been shown that different kinase sequences have different propensities to adopt DFG-out in the absence of inhibitor^9,10^, and the DFG-out back pocket has been suggested to have a lesser degree of sequence and structural homology between kinases^11^. However, the notion that type-II inhibitors developed to-date are more selective than type-I inhibitors has been brought into question^12,13^, suggesting that further consideration of the energetic contributions described above is required.

In order to fully exploit the target-selective potential of type-II inhibitors it is necessary to understand the underlying sequence-dependent principles that control the conformational preferences of their kinase targets, and the extent to which this has been diversified by evolution. This can, in principle, be directly approached using free-energy simulations to estimate the reorganization free-energy required for different kinases to adopt DFG-out, although this is computationally very expensive and of uncertain reliability for conformational changes involving large-scale loop reorganizations, such as the ∼17 Å “folding” of the activation loop that accompanies the transition from active DFG-in to the inactive, classical DFG-out state. To accommodate this limitation, we employ modern sequence-based computational methods to characterize the conformational selection process over the entire kinome, and combine the sequence-based results with structure-based free energy simulations with the goal of identifying evolutionarily divergent features of the energy landscape that control the preference of individual kinases for the active (DFG-in) vs inactive (DFG-out “folded activation loop”) states. To this end, we report evidence that TK catalytic domains have a molecular evolutionary bias that shifts their conformational equilibrium towards the inactive “folded activation loop” DFG-out state in the absence of activation signals. In contrast, STKs as a class have a more stable active conformation which is favored over the DFG-out state due to sequence constraints in the absence of other signals.

As described below, our analysis of a previously published kinome-wide assay suggests that TKs have properties which privilege the binding of type-II inhibitors in comparison to STKs, which leads us to hypothesize an evolutionary divergence in their conformational energy landscapes. To investigate this, we used a Potts Hamiltonian statistical energy model derived from residue-residue covariation in a Multiple Sequence Alignment (MSA) of protein kinase sequences to probe their conformational equilibria as previously described^9^. Using an approach that involves “threading” a large number of kinase sequences onto ensembles of DFG-in and DFG-out structures from the PDB and scoring them using the Potts Hamiltonian, we are able to view the evolutionary divergence in TK and STK conformational landscapes. To validate our results, we used the Potts statistical energy threading calculations to guide target selection for a set of more computationally intensive free-energy simulations. These simulations use type-II inhibitors as tools to probe kinase targets that have already reorganized to DFG-out, allowing us to estimate the free-energy of reorganization Δ*G*_*reorg*_ as the excess between the absolute binding free-energy calculated from simulations and the standard binding free-energy measured experimentally *in vitro*, which already includes the cost to reorganize. Although our methods avoid sampling the conformational change directly, we show how important structural determinants of the conformational change can be identified by analyzing residue-pair contributions to the Potts threading calculations, enabling us to reason about the molecular evolutionary basis for the differences in conformational behavior observed for TKs and STKs.

## Results

### Insights into the sequence-dependent conformational free-energy landscape

The binding of type-II inhibitors is achieved once a protein kinase has reorganized to the DFG-out with activation loop folded conformation (classical DFG-out). We sought initial insight into the conformational equilibrium from type-II binding data available publicly in the form of literature-reported dissociation constants (Kd). From the binding assay reported by Davis et al.^7^ we report a “hit” where an inhibitor binds to a kinase with K_d_ ≤ 10 μM. Using this criterion, a type-II inhibitor hit rate was calculated for each kinase. Analysis of the type-II hit rate distributions for STKs and TKs from the Davis assay (Fig. 1*A*) indicates that STKs, on average, have an unfavorable contribution to the binding of type-II inhibitors relative to TKs.

**Fig. 1.**
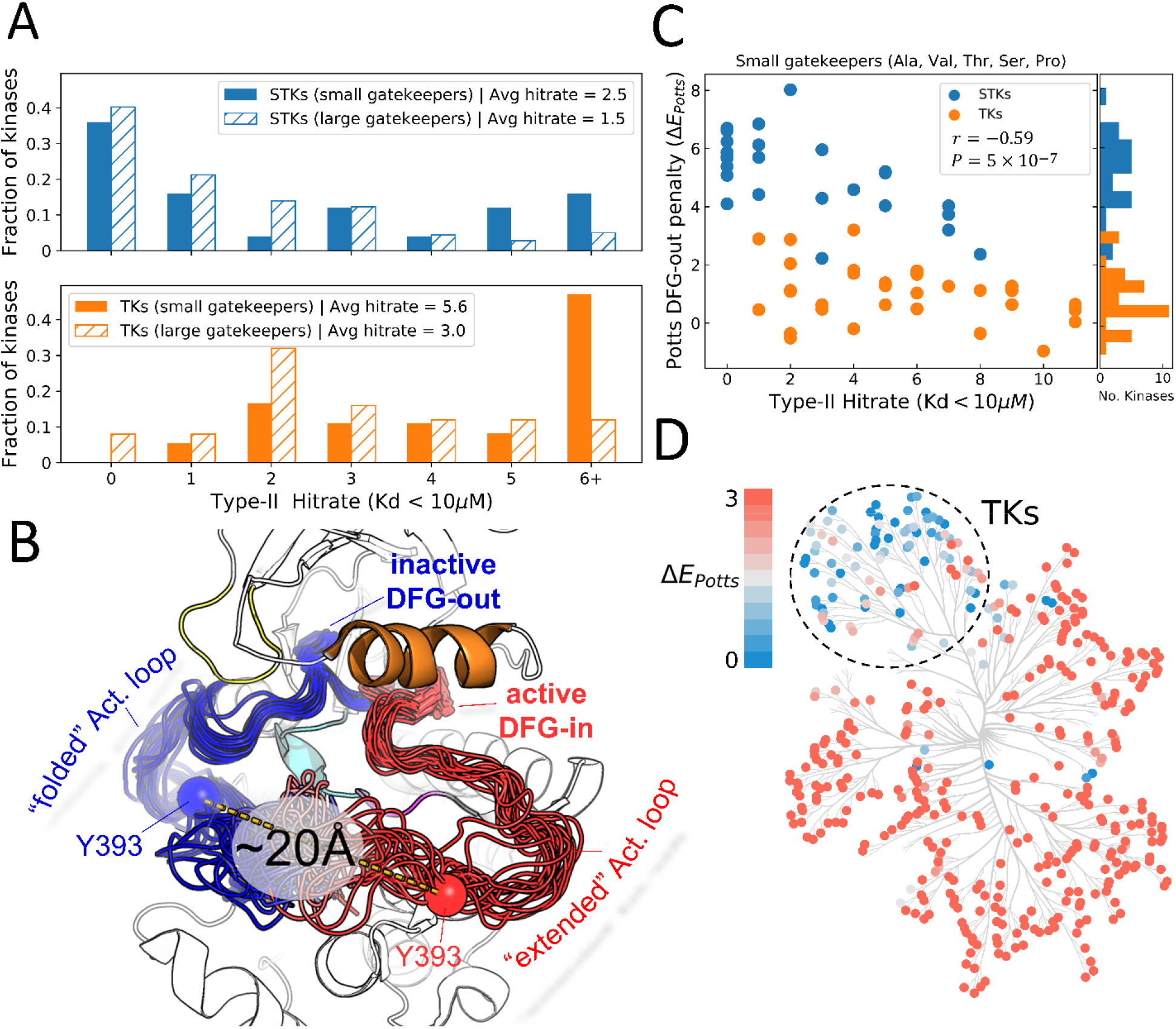
Viewing the conformational landscape of the human kinome. (A) Hit rate distributions from kinome-wide experimental binding assays with type-II inhibitors for human serine/threonine kinases (blue, top) and tyrosine kinases (orange, bottom) with small gatekeepers (solid bars) (sidechain volume < 110 Å^3^) and large gatekeepers (hatched bars) (sidechain volume > 110 Å^3^). (B) PyMol^14^ visualization of two conformational ensembles populated by Abl kinase from recent solution NMR experiments^15^. The active DFG-in conformation where the activation loop is “extended” (red, PDB: 6XR6) and the inactive classical DFG-out conformation where the activation loop is “folded” (blue, PDB: 6XRG) both exist in the absence of ligands, but there is a free-energy cost to transition between them.^15^ Type-II inhibitors preferentially bind to this folded DFG-out state. (C) Correlation between Potts DFG-out penalty (ΔE Potts) and hit rates for kinases with small gatekeepers only, to control for gatekeeper effects (Pearson correlation of −0.59, P < 0.001). (D) Potts DFG-out penalties calculated for the human kinome and plotted using CORAL^16^; the TK branch appears to have lower penalties relative to the rest of the kinome, which represent STKs.

The size of the gatekeeper residue is important for type-II binding as it controls access to a hydrophobic pocket adjacent to the ATP binding site that is traversed by type-II inhibitors^17–21^ and the size of the gatekeeper residue is thought to negatively affect type-II binding^17,22–24^. Because TKs tend to have small gatekeepers in comparison to STKs^17,25^ we considered this as a possible explanation behind the bias for TKs to have larger type-II hit rates. By plotting the hit rate distributions for STKs and TKs where the gatekeeper is either small or large (Fig. 1*A*) we confirm that gatekeeper size has an important influence on type-II binding for both STKs and TKs^22^. However, the hit rate distribution for TKs appears more sensitive to gatekeeper size than STKs. Even with small gatekeepers, there is a significant fraction of STKs that have hit rates of zero compared with TKs, suggesting the difference in hit rates between TKs and STKs cannot be accounted for primarily by the size of the gatekeeper residue.

Recent solution NMR experiments with Abl kinase revealed two DFG-out conformational states^15^; one where the DFG motif has flipped “DFG-in to DFG-out” but the activation loop remains in a “minimally perturbed” active-like conformation, and the other state is a classical “folded” DFG-out conformation where the activation loop has moved ∼17Å away from the active conformation (Fig. 1*B*) and the DFG motif is in a “classical DFG-out”^6^ or “BBAminus”^26^ state. Type-II inhibitors were shown to preferentially bind to this folded DFG-out state, confirming observations that Abl is almost always co-crystallized with type-II inhibitors in this conformation. This binding behavior is also exhibited by other kinases, suggested by the large number of activation loop folded DFG-out states seen in type-II bound co-crystal structures (Fig. S1). Hence, the importance of large-scale activation loop conformational changes in type-II binding and the large number of residue-residue contact changes involved in this transition^9^ (Fig. S2) suggests the sequence variation of the activation loop and the catalytic loop with which it interacts, might contour the conformational landscape differently for TKs compared with STKs. To investigate this, we used a Potts statistical energy model of sequence covariation to estimate the energetic cost of the active DFG-in (activation loop extended) ⟶ inactive DFG-out (activation loop folded) transition for human TKs and STKs (see *Methods*).

Patterns of coevolution of amino acids at different positions in an MSA are thought to largely reflect fitness constraints for fold stability and function between residues close in 3D space^27–30^, and these coevolutionary interactions can be successfully modeled by a Potts Hamiltonian^31,32^ which we inferred using Mi3-GPU, an algorithm designed to solve “Inverse Ising” problems for protein sequences with high accuracy^33^. The pairwise interactions from the Potts model can be used as a simple threaded energy function to estimate energetic differences between two conformations, based on changes in residue-residue contacts in the PDB^9^. We have calculated the threading penalty for all kinases in the human kinome. Our calculations show the Potts predicted DFG-out penalty (Δ*E*_*Potts*_), which is dominated by large-scale reorganization of the activation loop to the folded DFG-out state, is correlated with type-II hit rates (Fig. 1*C*) when controlling for gatekeeper size. From this, we determine that sequence variation of the activation loop and the contacts broken/formed by its large scale conformational change (Fig. S2) makes an important contribution to the binding affinity of type-II inhibitors.

Notably, our calculations over the entire human kinome show that the large majority of kinases with large Δ*E*_*Potts*_ (unfavorable conformational penalties) are STKs and the large majority of low-penalty kinases are TKs (Fig. 1*D*). To validate this finding, we next perform an independent and more computationally intensive prediction of the conformational reorganization energy of TKs and STKs for select kinase targets, chosen based on the kinome calculations of Δ*E*_*Potts*_ and type-II hit rates shown in Fig. 1, in which we use type-II inhibitors as probes in absolute binding free-energy simulations as described in the following section. By comparing the conformational penalties predicted from these structure-based molecular dynamics free-energy simulations with the Potts conformational penalty scores, we also identify the scale of Δ*E*_*Potts*_ in physical free-energy units. This allows us to predict physical conformational free energies based on Potts calculations which can be carried out at scale on large numbers of sequences, to evaluate the evolutionary divergence of the conformational penalty between STKs and TKs generally.

### Structure-based free-energy simulations guided by the sequence-based Potts model

Relative binding free-energy simulations are now widely employed to screen potent inhibitors in large-scale drug discovery studies^34^. These methods are used to determine the relative free-energy of binding between ligands that differ by small substitutions, which permits one to simulate along an alchemical pathway that mutates one ligand to another. By leaving the common core scaffold unperturbed, the cost and difficulty of sampling the transition between unbound (apo) and bound (holo) states of the system are avoided^34–36^. Alternatively, alchemical methods to determine absolute binding free-energy (ABFEs), such as the “double decoupling” method employed in this work, sample the apo ⟶ holo transition along a pathway that decouples the entire ligand from its environment. While more computationally expensive, the advantage of ABFE is that the computed Δ*G*_*bind*_ can be directly compared with experimental binding affinities, and successful convergence does not rely on the structural similarity of compounds being simulated^37–43^.

Our alchemical ABFE simulations of type-II inhibitors binding to TKs and STKs simulate the apo and holo states of the kinase domains in the classical DFG-out conformation with the activation loop folded, starting from the experimentally determined co-crystal structure of the holo state. The apo state remains DFG-out with the activation loop folded throughout the simulations, and therefore the calculated absolute binding free-energy 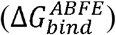 excludes the cost to reorganize from DFG-in (Δ*G*_*reorg*_). On the other hand, standard binding free-energies 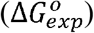 determined experimentally from inhibition or dissociation constants (Eq. 1) implicitly include the free-energy cost to reorganize. Therefore given the experimentally determined total binding free energy 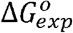, ABFE simulations can be used to separate the free-energy of ligand-receptor association in the inactive state 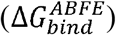 from the cost to reorganize from the active to inactive state Δ*G*_*reorg*_ (Eq. 3)^44–46^.

We calculated 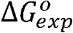 (Eq. 1) from literature reported IC_50_ or *K*_*d*_ values, where the standard concentration *C*_0_ is set to 1 *M*

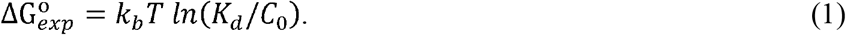

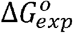 can be expressed as the sum of the free energy change to reorganize from the active to inactive state, Δ*G*_*reorg*_ plus the free energy to bind to the inactive state 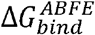 (Eq. 2). Δ*G*_*reorg*_ is therefore the excess free-energy difference between 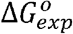 and 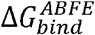 (Eq. 3)

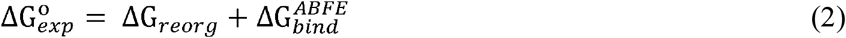

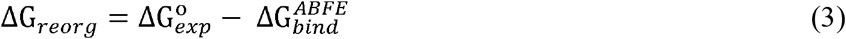

Type-II inhibitors generally bind when the activation loop is in a folded DFG-out conformation (Fig. 1*B*)., which presents major challenges for direct simulations to determine the free energy cost of the conformational change in contrast to the method employed here (Eq. 3).

Because the type-II inhibitor imatinib is co-crystallized in a type-II binding mode with MAPK14 (p38*α*), an STK, and several other TKs (e.g. ABL1, DDR1, LCK, CSF1R, KIT and PDGFRA), we chose this inhibitor as an initial probe of our hypothesis that TKs evolved to have lower Δ*G*_*reorg*_ than STKs (Fig. 2). In this example we note that TKs bind strongly to imatinib (“STI” in Fig. 2) with an average 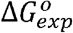 of −9.3 kcal/mol, in contrast to the STK MAPK14 which binds this drug very weakly (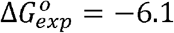 kcal/mol). At face-value this appears consistent with our analysis from Fig. 1, where we calculated a large Potts DFG-out penalty for MAPK14 (Δ*E*_*Potts*_ = 5.2) and low penalties for TKs, suggesting that the weak binding of imatinib to MAPK14 is due at least partially to large Δ*G*_*reorg*_. To confirm this, we used ABFE simulations with the imatinib: MAPK14 complex to evaluate Eq. 3, confirming that MAPK14 incurs a large penalty to adopt the DFG-out conformation with the activation loop folded (Δ*G*_*reorg*_ = 5 kcal/mol) (Fig. 2).

**Fig. 2.**
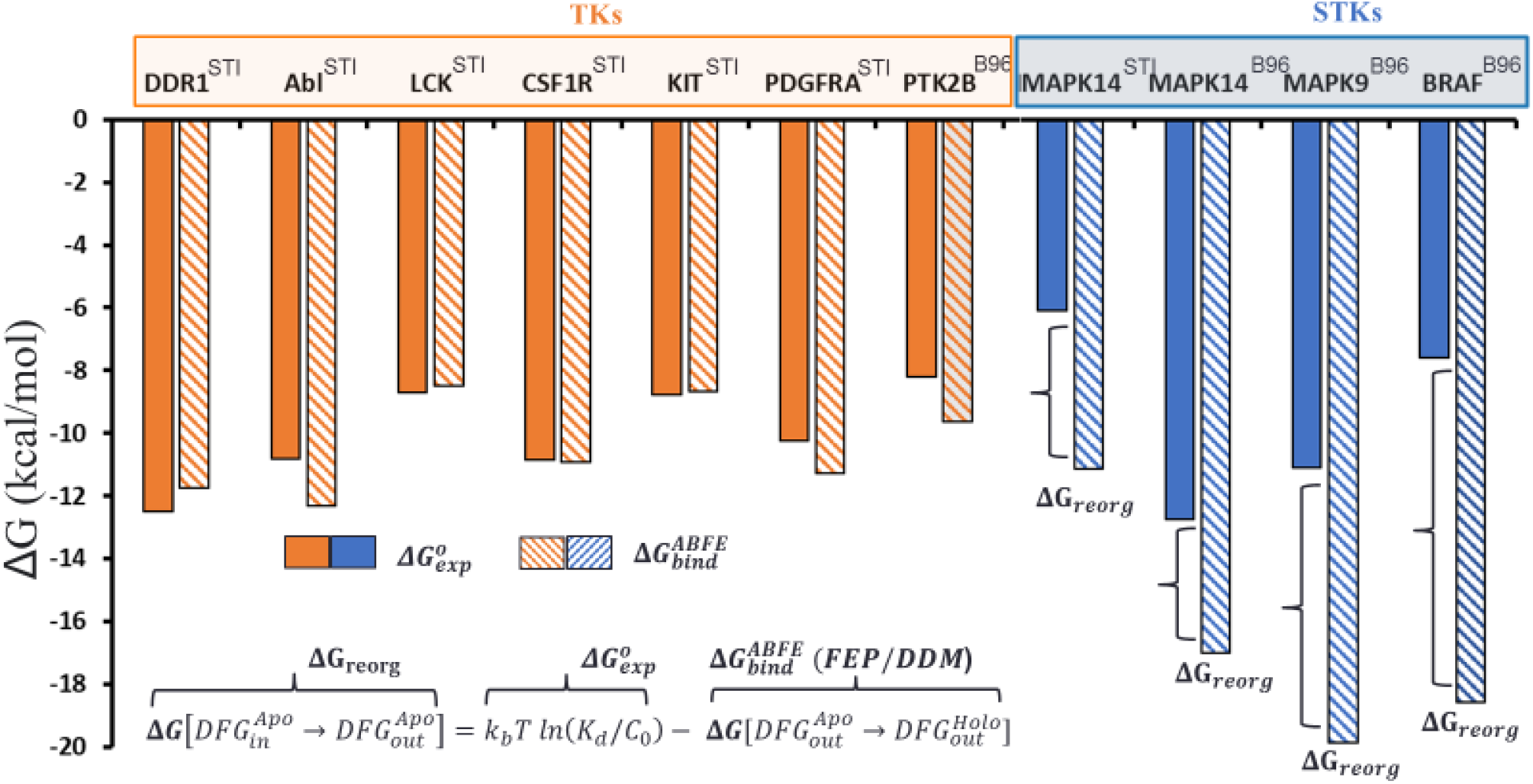
Overview of the conformational landscapes between STKs and TKs from ABFE simulations, where we compare ΔG_bind_ (hatched bars) from binding free-energy simulations with ΔG_exp_ (solid bars) for the type-II inhibitors imatinib (PDB code: STI) and BIRB-976 (PDB code: B96) vs several TKs (orange) and STKs (blue).

Despite the large Δ*G*_*reorg*_ predicted for MAPK14 by both the Potts model and the simulations with imatinib described above, highly potent type-II inhibitors have been successfully developed for this kinase. For example, BIRB-796^47^ binds to MAPK14 about 7 kcal/mol more strongly than imatinib. This stronger binding of BIRB-796 is captured by 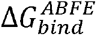 from our simulations (Fig. 2) and the calculated value of Δ*G*_*reorg*_ for this complex (Δ*G*_*reorg*_ ≈ 4kcal/mol) is very close to the corresponding estimate of Δ*G*_*reorg*_ based on simulations with imatinib (Fig. 2). Importantly, this result suggests that STKs can be potently inhibited by type-II inhibitors despite their large Δ*G*_*reorg*_. To support this, we performed additional ABFE simulations with BIRB-796 and calculated Δ*G*_*reorg*_ for two additional STKs predicted to have large reorganization penalties (MAPK9 and BRAF, Δ*E*_*Potts*_ ≥ 4). We calculated Δ*G*_*reorg*_ > 8 kcal/mol for MAPK9 and BRAF, which is consistent with predictions from the Potts model, and comparison of 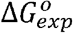 and 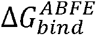 in Fig. 2 confirms that BIRB-796 is able to overcome the large Δ*G*_*reorg*_ of certain kinases to attain high experimental potencies (e.g. MAPK14 and MAPK9) (Fig. 2). To further validate this result, we calculated Δ*G*_*reorg*_ via ABFE simulations of BIRB-796 binding to a TK predicted by the Potts model to have a low penalty (PTK2B, Δ*E*_*Potts*_ < 1), which again shows consistency with our Potts prediction of the conformational landscape (Fig 2). The relatively weak value of 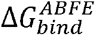 for this kinase compared with MAPK14 is also consistent with observations of the BIRB-796: PTK2B co-crystal structure (PDB: 3FZS), where the binding mode in PTK2B is more weakly associated with the ATP pocket in comparison with MAPK14^48^.

The analysis above provides initial support for our hypothesis about the evolutionarily divergent STK and TK conformational landscapes. To further develop this approach, we identified five STKs and five TKs which are predicted by the Potts threading calculations to have large and small Δ*G*_*reorg*_ respectively, and for which there is sufficient experimental structural and inhibitory data (co-crystal structures and binding constants) to calculate an average Δ*G*_*reorg*_via Eq. 3 for each target using multiple type-II inhibitor probes. For each of the TK and STK targets, these sets of calculations can be visualized as a linear regression of 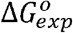 vs 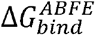 where the slope is constrained to one, consistent with Eq. 2 (see SI for details). We employed this workflow for the set of five TK and five STK targets by simulating 22 and 23 type-II inhibitor complexes, respectively.

The result of this workflow for the set of five TKs and their type-II complexes revealed a low average Δ*G*_*reorg*_ of < 1 kcal/mol (Fig. 3*B*), consistent with our initial predictions from Potts Δ*E*s and type-II hit rates (Table 1). On the other hand, the binding free energy simulations for the set of five STKs and their type-II complexes show an average of ∼6 kcal/mol of Δ*G*_*reorg*_ is required for these kinases to adopt DFG-out conformation, which is also consistent with our initial predictions from the Potts model (Table 1). To verify that the large Δ*G*_*reorg*_ identified for STKs is a property of conformational selection for DFG-out rather than systematic overestimation of 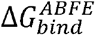 for these kinases, we performed ABFE simulations of type-I inhibitors binding to the same set of STKs (an additional 23 complexes). For the binding of type-I inhibitors, we expect there to be no reorganization penalty due to the lack of DFG-out conformational selection in their binding mechanism. As anticipated, the calculated values of 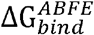 for type-I inhibitors are very close to their experimental binding affinities 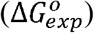 (Fig. 3*A*).

**Fig. 3.**
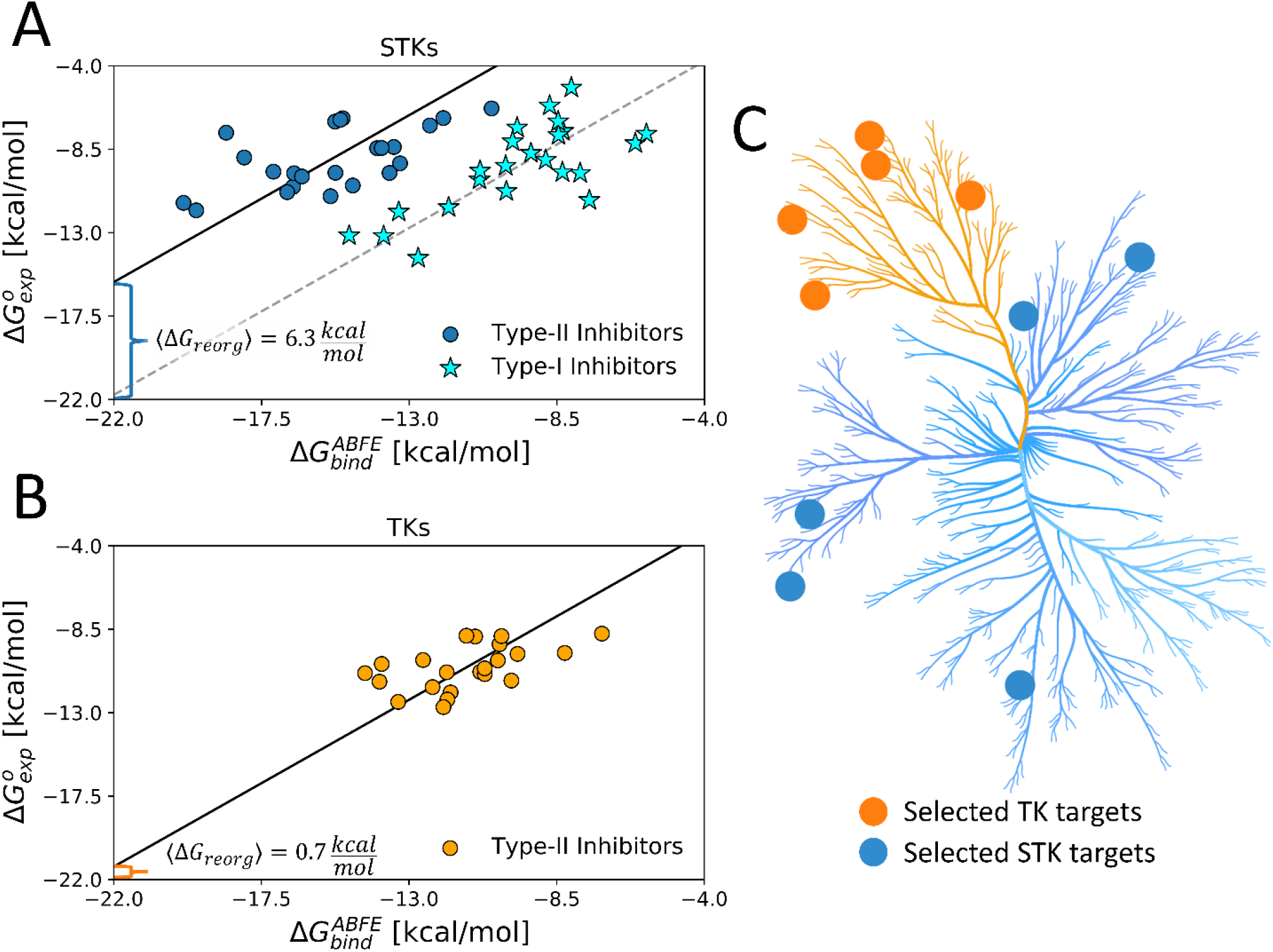
(A) The average ΔG_reorg_ calculated via absolute binding free energy simulations with 23 type-I (stars) and 23 type-II inhibitors (circles) complexes in the active DFG-in and inactive DFG-out state, respectively, computed from five STK targets (Table 1) and (B) computed with 22 type-II inhibitors vs five TK targets in the DFG-out state (Table 1). (C) Kinome plot created with CORAL^16^ illustrating the selection of five TKs and five STKs which are detailed in Table 1.

**Table 1.** Type-II hit rates from Davis et al. and Potts threaded energy penalties from Fig. 1 were used to guide the selection of five STK and five TK targets for absolute binding free-energy simulations. For some kinases, the hit rate binary classifier captures a set of relatively weak hits with average binding which, in context with large Potts penalty (see Fig. 1*D*), might be explained by a large ΔG_reorg_ incurred for the folded DFG-out state (Fig. 1*B*).

| Kinase | Class | Hit rate<sup>a</sup><br>($K_d < 10 \mu M$) | Potts Penalty<sup>b</sup><br>($\Delta E_{Potts}$) | Calculated<br>$\Delta G_{reorg}^c$ |
| --- | --- | --- | --- | --- |
| MELK | STK | 3 | 5.9 | $5.6 \pm 0.2$ |
| MAPK9 | STK | 5 | 4.7 | $6.9 \pm 0.3$ |
| CDK2 | STK | 2 | 5.3 | $7.7 \pm 0.2$ |
| IRAK4 | STK | 0 | 2.5 | $5.4 \pm 0.2$ |
| BRAF | STK | 7 | 4.0 | $6.5 \pm 0.1$ |
| ABL1 | TK | 10 | -1.0 | $1.3 \pm 0.3$ |
| LCK | TK | 11 | 0.5 | $1.0 \pm 0.3$ |
| TIE2 | TK | 6 | 1.1 | $-0.3 \pm 0.2$ |
| NTRK2 | TK | 6 | -1.0 | $1.7 \pm 0.1^d$ |
| DDR1 | TK | 11 | 0.4 | $0.3 \pm 0.3$ |

<sup>a</sup> Type-II inhibitors only, data from ref. 7.
<sup>b</sup> Calculations from Fig. 1d.
<sup>c</sup> $\Delta G_{\text{reorg}}$ was calculated from Equation 3. Reported standard deviations were calculated by propagating error from the simulations used in the calculation of average $\Delta G_{\text{reorg}}$ in units of kcal/mol (see Supplementary for statistics from individual simulations).
<sup>d</sup> Average and standard deviation calculated from two simulated complexes.

We find that the set of type-II inhibitors complexed with STKs in this dataset tend to have more favorable binding free energies to the reorganized receptor 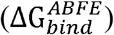 than type-II inhibitors complexed with TKs, as shown by their distribution along the horizontal axes in Fig. 3. The reason for this can be understood as a consequence of selection bias. Our selection of STK complexes for this study usually involved lead compounds from the literature, which were designed for high on-target experimental potency and published for their pharmaceutical potential, similar to BIRB-796: MAPK14 which is a tightly bound complex with high experimental affinity despite the large Δ*G*_*reorg*_ incurred by this kinase (Fig. 2). This tight binding is reflected by the favorability of the 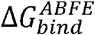 term which must be implicitly tuned by medicinal chemists to overcome the large Δ*G*_*reorg*_ found in STKs. Meanwhile, the chemical space of type-II inhibitors studied against TKs appears to be privileged by their low Δ*G*_*reorg*_, judging by the comparably weak 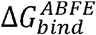 for these complexes. This ultimately gives rise to similar experimental potencies for the binding of type II inhibitors to TKs and STKs plotted in Fig. 3.

The results of the molecular dynamics binding free energy simulations when combined with experimental binding affinities, reveal significant differences in the conformational free-energy landscapes between STKs and TKs. The DFG-in (activation loop extended) to DFG-out (activation loop folded) reorganization penalties are strongly correlated with corresponding Δ*E*s calculated from the Potts model (*R*^2^ = 0.75, *P* ≈ 10^−3^) emphasizing the connection between coevolutionary statistical energies in sequence space and physical free-energies in protein conformational space (Fig. 4). From this relationship, we can approximate a scale for the Potts Δ*E* scores in physical free-energy units which describes the conformational landscapes of folded proteins in a similar manner to that of an earlier study of protein folding landscapes^49^; we find that a Potts statistical energy difference Δ*E* of one unit corresponds approximately to 1.3 kcal/mol of Δ*G*_*reorg*_.

**Fig. 4.**
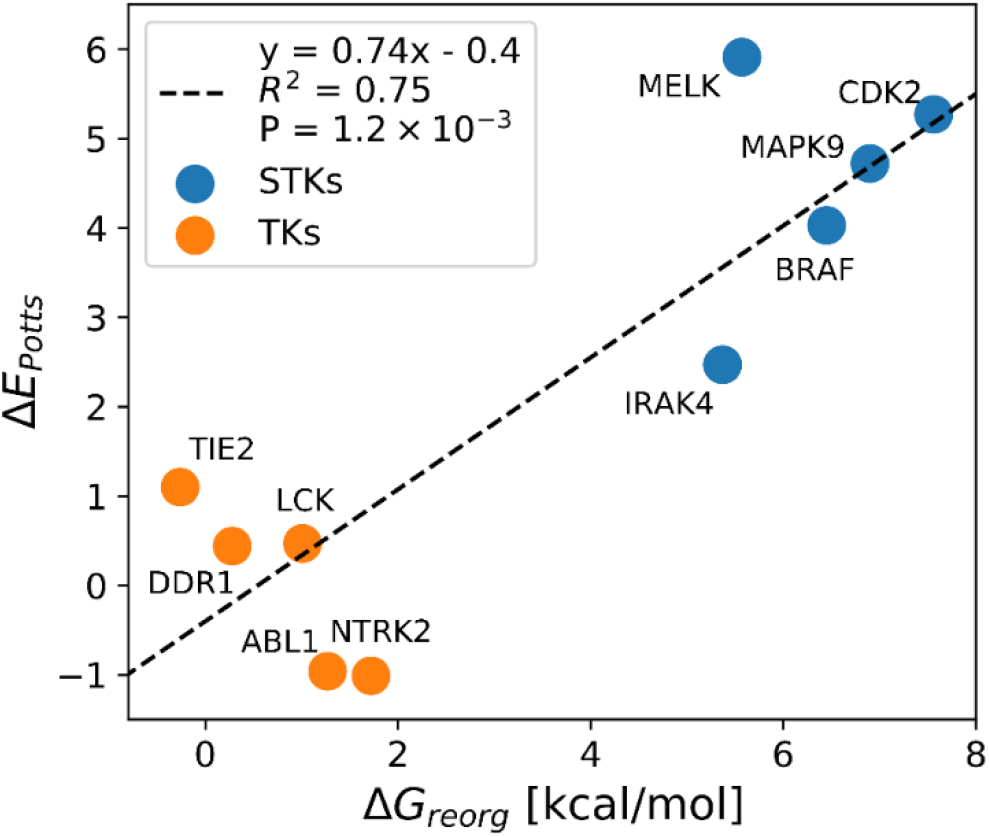
Correlation between ΔE_Potts_ and averaged calculations of ΔG_reorg_ for five TKs and five STKs (Table 1).

### Structural and evolutionary basis for the divergent TK and STK conformational landscapes

The consistency of the predictions between Δ*G*_*reorg*_ and Δ*E*_*Potts*_ identified from Potts-guided free-energy simulations (Fig. 4) led us to investigate whether the observed differences in the free energy to reorganize from the active to inactive state is a more general feature that distinguishes TKs from STKs. To this end, we extracted ∼200,000 STKs and ∼10,000 TKs from the large MSA of Pfam sequences used in the construction of our Potts model, based on patterns of sequence conservation that clearly distinguish the two classes (see *Methods*). For each sequence, we calculated Δ*E*_*Potts*_ threaded over the structural database (a total of 4268 active DFG-in and 510 classical DFG-out PDB structures) and plotted the distributions for TKs and STKs, revealing a bias for STKs towards larger Potts conformational penalties (Fig. 5). The average difference between these distributions, ΔΔ*E* = 3.2, is extremely unlikely to be observed by chance (*P* ≤ 10^−15^, see *Methods)* and supports the hypothesis that TKs are evolutionarily biased towards a lower free-energy cost to adopt the classical “folded activation loop” DFG-out conformation (Δ*G*_*reorg*_) compared to STKs. We estimate that ΔΔ*E* = 3.2 corresponds to ∼ 4.3 kcal/mol based on the analysis summarized in Fig. 4.

**Fig. 5.**
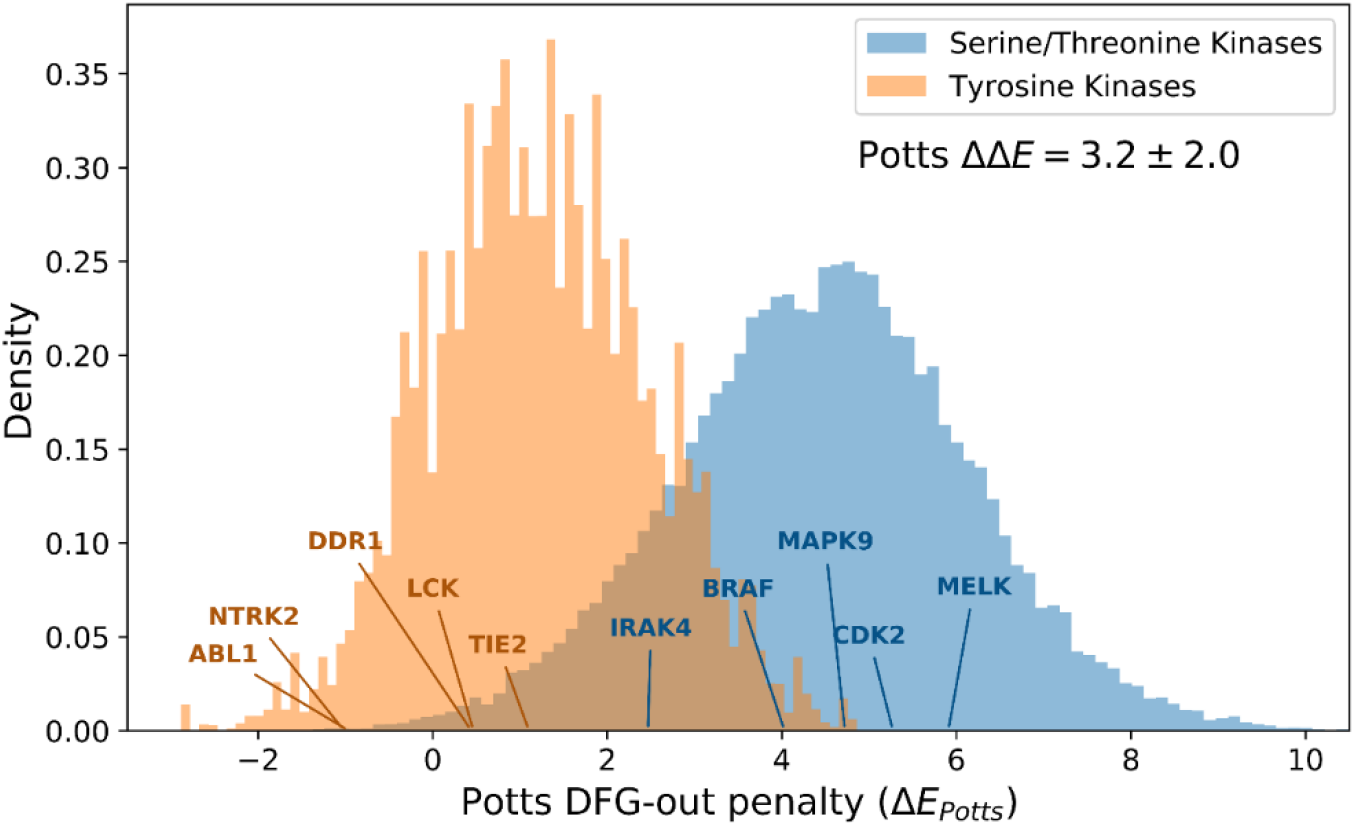
The distributions of Potts conformational penalties for (orange) 10,345 Tyrosine Kinases and (blue) 210,862 Serine/Threonine Kinases show that Tyrosine Kinases tend to have smaller energetic penalties on average. The difference between in averages between these distributions is shown (ΔΔ*E* = 3.2), which we estimate to be ≈ 4.3 kcal/mol based on the analysis in Fig. 4).

To gain insight into the molecular basis for this effect which distinguishes the conformational landscape of TKs from STKs, we examined the residue-residue interactions that make the most significant contributions to the observed ΔΔ*E*. The difference between average Potts threaded-energy penalties, ΔΔ*E* = ⟨Δ*E*⟩_*STKs*_ − ⟨Δ*E*⟩_*TKs*_, can also be written as a sum over pairs of alignment positions *i* and *j* along length *L* of the aligned kinase domains, 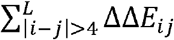 (see *Methods* for details). We find that ∼75% of the total contribution to ΔΔ*E* (approximately 3 kcal/mol) can be traced back to a small number (ten) of residue-residue interactions involving the activation loop, suggesting that mutations within the activation loop are largely responsible for the evolutionary divergence between the conformational free-energy landscapes of TKs and STKs. These interactions occur between important structural motifs responsible for controlling the stability of the active “extended” conformation of the activation loop (Fig. 6*A*), especially the activation loop N-terminal and C-terminal “anchors”^50^, and the regulatory “RD-pocket” formed by the H**RD** motif of the catalytic loop which functions to stabilize or destabilize this conformation depending on the activation loop’s phosphorylation state^50,51^. The remaining top ΔΔ*E*_*ij*_ correspond to contacts that stabilize the DFG-out “folded” conformation of the activation loop for TKs, wherein the kinase’s substrate binding site recognizes its own activation loop tyrosine (Fig. 6*C* *right*)^52^. We describe these interactions below, focusing on the strongest effects involving these structural motifs that lead to differences in the conformational free-energy landscapes of TKs and STKs. The residue nomenclature we use in our descriptions follows the format 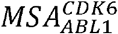, which is the unique residue numbering in our MSA followed by Abl1 (TK) numbering in the subscript (active PDB: 2GQG, inactive PDB: 1IEP) and CDK6 (STK) numbering in the superscript (active PDB: 1XO2, inactive PDB: 1G3N), corresponding to the original PDB files used to generate Fig. 6*B* and Fig. 6*C*.

**Fig. 6.**
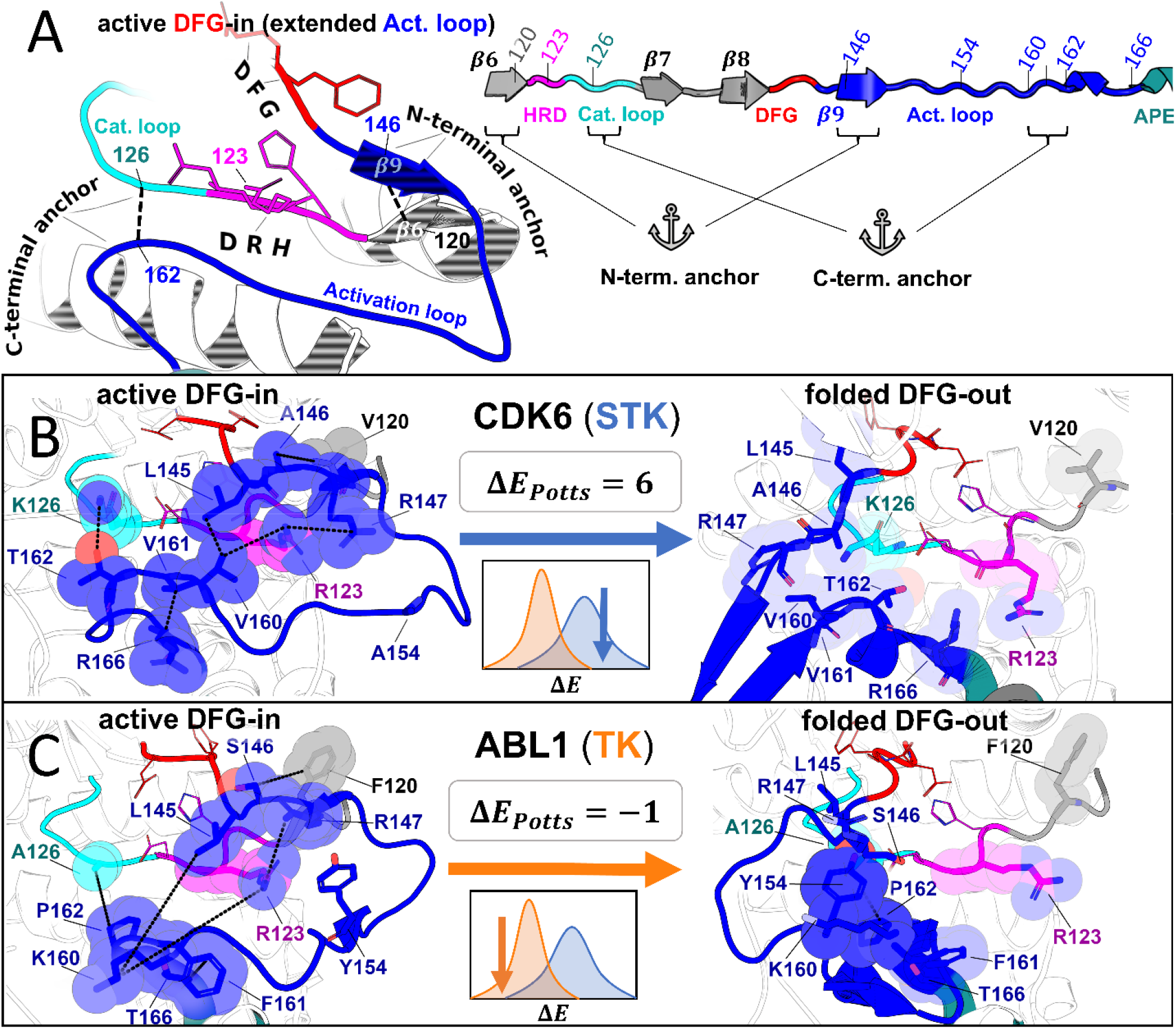
Molecular basis for the evolutionary divergence between TKs and STKs, based on the top interaction pairs that contribute to the result in Fig. 5 which are discussed in the main text. (A) Diagrams depicting general features of the active “extended” conformation of the activation loop (left) and the primary structure of these motifs in our MSA (right) with the HRD (pink), DFG (red), and APE (teal) motifs color-coded for reference. (B-C) Structural examples of a representative STK (CDK6) and TK (Abl) in the active DFG-in conformation (left) and the folded DFG-out conformation (right). Residues are labeled according to their position in our MSA, and colored according to *A*. The inset (center) displays the ΔE_Potts_ of the reference kinase derived from Fig. 5 as well as a cartoon depicting their location in the distributions. The diagrams of CDK6 in the active DFG-in conformation (PDB: 1XO2, chain B), CDK6 in the folded DFG-out conformation (PDB: 1G3N, chain A), Abl in the active DFG-in conformation (PDB: 2G2I, chain A), and Abl in the folded DFG-out conformation (PDB: 1IEP, chain A) were generated with PyMol.^14^ All ligands and some backbone atoms were hidden for clarity.

When the activation loop is in an active “extended” conformation, the N-terminal anchor^50^ is formed by a *β*6 strand in the N-terminal of the activation loop (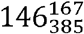 to 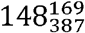) and a *β*9 strand near the N-terminal of the catalytic loop (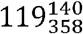 to 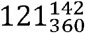). In the inactive “folded” conformation of the activation loop, the N-terminal anchor is completely broken as the *β*6 and *β*9 strands are peeled apart (Fig. 6*B* and Fig. 6*C*). The Potts model suggests this conformational change is energetically penalized in STKs due to favorable contacts between position 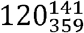 in the *β*6 strand and 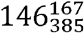 in the *β*9 strand; in 20% of STKs we observe *V*120 and *A*146 at these positions (Fig. 6*B*). These residues have a significantly more favorable Potts interaction energy than the TK-prevalent pair (*F*120, *S*146) which appears in 17% of TKs and only 1% of STKs (Fig. 6*C*). As reported previously, position 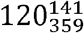 in TKs is a frequent site of drug resistance mutations to type-II inhibitors^53,54^, particularly the F120I mutation (F359I in Abl numbering). This mutation in Abl changes the interaction pair from (*F*120, *S*146), seen in only 1% of STKs, to (*I*120, *S*146) which is seen in 15% of STKs. The mutant “STK-like” interaction pair (*I*120, *S*146) is energetically more favorable according to the Potts model, suggesting that part of the Abl type-II inhibitor resistance mechanism of F120I (F359I) involves increasing the free-energy penalty for the active DFG-in / extended activation loop ⟶ classical DFG-out / folded activation loop transition. Consistent with this view, it was found that it is possible to treat patients with this drug-resistance mutation using dasatinib^55,56^: a type-I inhibitor that does not require access to the DFG-out conformation.

The “RD-pocket”^50^ is a conserved basic pocket formed by the Arg and Asp residues of the HRD motif ^51^ (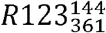 and 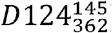) and a positively charged Lys or Arg that is often present in the N-terminal anchor of the activation loop 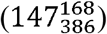. 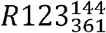 in the HRD motif and Lys/Arg 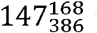 in the N-terminal anchor form an unfavorable like-charge interaction when the activation loop is in the active, extended conformation^50^. Kinase activation is typically a complex process involving many layers of regulation from other protein domains, cofactors, and phosphorylation events^57^; however, a general activation mechanism that applies to the majority of kinases involves quenching the net-charge of the RD-pocket by addition of a negatively charged phosphate group to a nearby residue in the activation loop, stabilizing the active conformation. The conservation of this regulatory mechanism in most protein kinases, particularly those bearing the HRD-Arg residue (termed “RD-kinases”^51^), explains why Lys or Arg is frequently observed at position 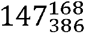 of the N-terminal anchor of the activation loop. However, RD-TKs prefer Arg at this position (78%) which the Potts model suggests has a greater destabilizing effect on the active conformation than Lys (9%) due to interactions with HRD-Arg. The activation loop Arg also forms part of an electrostatic interaction network that stabilizes the “Src-like inactive” conformation in TKs^58,59^, a conformation with a “partially” folded activation loop (Fig. S1) that is suggested to be an intermediate state along the transition to DFG-out^60,61^. On the other hand, RD-STKs display *K*147 more frequently (26%), which the Potts model suggests interacts more favorably with the HRD-Arg independently of activation loop phosphorylation, thus contributing to greater stabilization of the active DFG-in conformation for RD-STKs in comparison with TKs. Additionally, the Potts statistical energy analysis suggests that packing interactions between HRD-Arg and 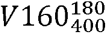 located near the activation loop C-terminal also contributes to phosphorylation-independent stabilization of the active conformation in RD-STKs (Fig. 6*B* *left*), appearing in 28% of RD-STKs and only 2% of RD-TKs. In RD-TKs, the residue at this position is usually Arg or Lys which appears to be repelled from the RD-pocket (Fig. 6*C* *left*) and is suggested by the Potts model to result in a less stable active conformation.

The largest ΔΔ*E*_*ij*_ term, which contributes 16% of the total difference in average Potts conformational penalties between STKs and TKs, comes from an interaction pair in the C-terminal of the activation loop 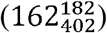 and the C-terminal of the catalytic loop 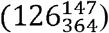. These residues form part of the “C-terminal anchor”^50^ which is important for creating a suitable binding site for the substrate peptide. The C-terminal anchor residue 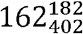 is Pro in TKs and typically Ser or Thr in STKs^62^. In STKs, the sidechain hydroxyl of this residue forms a hydrogen bond with 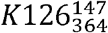 in the catalytic loop, creating a stable binding site for substrate phosphoacceptor residues and stabilizing the C-terminal anchor^62^. *K*126 is also directly involved in catalysis by interacting with and stabilizing the gamma phosphate of ATP^63^, hence it is often referred to as the “catalytic lysine”. The hydrogen bond between *K*126 and *S* or *T*162 is almost always formed in the active DFG-in conformation, and we observe breakage of this hydrogen bond in many STKs crystallized in the DFG-out / activation loop folded conformation (e.g. CDK6, Fig. 6*B*), suggesting that deformation of the C-terminal anchor contributes an energetic penalty for the active ⟶ inactive conformational change. In TKs however, the catalytic lysine is almost always mreplaced with Ala, with the exception of a few TKs (e.g. c-Src) which have instead adopted Arg at this position. The C-terminal anchor of TKs containing *A*126 and *P*162 is less stable in comparison to STKs containing (*K*126, *T*162) or (*K*126, *S*162) for which the Potts coupling is very favorable, consistent with the structural observation that (*A*126, *P*162) form weak interactions (Fig. 6*C*). Our analysis suggests the interaction pair (*A*126, *P*162) weakens the C-terminal anchor, leading to a less stable active conformation in TKs as compared with STKs. Another significant contribution to the stabilization of the C-terminal anchor in the active DFG-in conformation for STKs comes from interactions between the residue pair 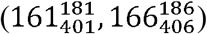 which are both located within the activation loop. We observe (*G*161, *M*166) at this position pair in 33% of STKs, but never in TKs (Fig. S4*D*). The Potts coupling between these residues is highly favorable. In contrast, we observe (*L*161, *M*166) in 40% of TKs but never in STKs (Fig. S4*H*), which have weaker coupling. The bulky sidechains of (*L*161, *M*166) observed in TKs causes the activation loop to “bulge” in this C-terminal region which has been previously identified as a feature of TKs that helps shape the substrate binding site to accommodate Tyr residues^50^. In addition to this paradigm, our analysis suggests that the C-terminal bulge results in weaker structural constraints on the active conformation relative to STKs.

In summary, TKs are suggested by the Potts statistical energy model which is based on sequence covariation, to have on average, weaker N-terminal anchor, RD-pocket, and C-terminal anchor interactions than STKs. This mechanism of shifting the TK conformational equilibrium away from the active DFG-in / extended activation loop conformation can explain seven of the top ten ΔΔ*E*_*ij*_, accounting for ∼80% of these contributions to the divergence between the STK and TK conformational landscapes. The remaining ∼20% of the top contributions can be attributed to residue-residue interactions that occur within the folded DFG-out conformation wherein the activation loop of TKs binds to the kinase’s own active site as though it were engaging a peptide substrate in trans (Fig. 6*C* *right*)^50^. STKs, however, rarely adopt a folded DFG-out conformation with this property, and instead the activation loop is typically found to be unresolved and/or projecting outwards towards solvent (Fig. 6*B* *right*). The Potts model suggests that this substrate mimicry of the folded DFG-out activation loop observed in TKs is highly dependent on the presence of a Tyr phosphorylation site at position 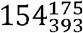 in the activation loop (Fig. 6*C*). In the active conformation, the anionic p*Y*154 stabilizes the active conformation by binding to the basic RD-pocket (Fig. 6*C* *left*, phosphate not shown). However, in the (unphosphorylated) folded DFG-out conformation, this *Y*154 mimics a substrate by stacking against the TK-conserved 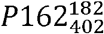 residue (Fig. 6*C* *right*)^50^. The substrate mimicking nature of this binding mode is demonstrated by the autophosphorylation dimer structure of tyrosine kinase FGFR3 (PDB: 6PNX) solved recently (Fig. S4*K*)^64^.

The striking connection between the ability of TKs to phosphorylate tyrosine substrates and their enhanced access to the DFG-out conformation via substrate-competitive contacts from their own activation loop described above suggests an evolutionary model for the TK conformational behavior characterized in this work. In this model, the coevolution of residues that form substrate-competitive contacts in folded DFG-out appears to be a byproduct of the evolutionary pressure for TKs to phosphorylate other TKs on their activation loop tyrosine residues. STKs, on the other hand, have optimized the binding of Ser and Thr substrates via a different binding mode^57^ which does not have the same energetic feedback with the stability of the folded DFG-out conformation. Additionally, the catalytic domains of TKs appear to have a less energetically stable “extended” activation loop conformation than STKs, which may have encouraged the evolution of more complex mechanisms of allosteric regulation and autophosphorylation which are highly important regulatory mechanisms in TKs^64–66^. The combined effect of these two TK phenotypes, the former favoring the *de*stabilization of folded DFG-out and the latter favoring *de*stabilization of active DFG-in, may explain their low free-energy cost for the DFG-in ⟶ DFG-out conformational change compared with STKs.

## Discussion

In this work, we have combined sequence and structure-based approaches to analyze the conformational free energy difference between active DFG-in and inactive DFG-out kinase states. Using a Potts statistical energy model derived from residue-residue covariation in a kinase family multiple sequence alignment, we first threaded all human Serine/Threonine Kinase sequences (STKs) and Tyrosine Kinase sequences (TKs) onto large ensembles of active DFG-in and classical DFG-out structures from the PDB. We found distinctly different distributions of threading scores for Serine/Threonine kinases compared with Tyrosine kinases, with STKs having a significant conformational reorganization penalty compared with TKs. The molecular basis for the evolutionary divergence in the conformational landscapes was analyzed; a substantial contribution to the difference is associated with sequence position pairs that couple the N and C terminal anchor residues of the activation loop to N and C terminal residues in the catalytic loop, according to the Potts statistical energy analysis. We then used the Potts statistical energy model to guide the selection for structure-based molecular dynamics binding free energy simulations of 68 protein-ligand complexes; using the calculated binding free energy estimates together with experimental values, we were able to estimate free-energy costs for the large-scale (∼17 - 20 Å) conformational change of the activation loop by an indirect approach. The structure-based estimates of the reorganization free energy penalties are consistent with the sequence-based estimates. Additionally, the strong correlation between Δ*G*_*reorg*_ and Δ*E*_*Potts*_ identified in this study reveals that the conformational landscape has a strong sequence dependence with STKs having a ∼4 kcal/mol conformational free energy bias favoring the active state over the inactive state relative to TKs (Fig. 4). We note that the most potent type-II inhibitors from the literature which target STKs bind with nanomolar K_d_s, similar to that for TKs, despite the substantial additional reorganization penalty that STKs must overcome. This suggests that medicinal chemists have implicitly been able to exploit particularly favorable characteristics of the type-II binding pocket to design inhibitors with extremely strong affinities to the DFG-out (activation loop folded) receptor conformations of STKs, and that further analysis of the molecular basis for this tight binding could provide a basis for designing more selective inhibitors.

## Materials and Methods

### Enumeration of absolute binding free-energy simulations

In this work, we have performed all-atom molecular dynamics simulations in explicit solvent for a total of 94 different kinase-inhibitor complexes to calculate absolute binding free-energies (ABFEs) via the alchemical double decoupling method. 74 of these free-energy calculations were guided by insights from the Potts model, specifically the patterns of Potts conformational penalties plotted in Fig. 1D. 68 of these complexes correspond to the ABFEs plotted in Fig. 3, of which 23 type-II inhibitors and 23 type-I inhibitors ABFEs are plotted for STKs in Fig. 3*A*, and 22 type-II inhibitors ABFEs for TKs are plotted in Fig. 3*B*. Six additional Potts-guided ABFEs corresponding to CSF1R, KIT, PDGFRA, MAPK14, and PTK2B are included in Fig. 2. An additional 20 type-I and type-I ½ ABFEs were calculated as part of our benchmarking procedure described in the Supplementary Information.

### Multiple Sequence Alignment (MSA) and Classification of Serine/Threonine vs Tyrosine Kinases

An MSA of 236,572 protein kinase catalytic domains with 259 columns was constructed as previously described.^67^ STKs and TKs were classified based on patterns of sequence conservation previously identified by Taylor and co-workers^62^; characteristic sequence features of TKs and STKs which form their respective phosphoacceptor binding pockets are found at the HRD+2 (Ala or Arg in TKs, Lys in STKs) and HRD+4 (Arg in TKs, variable in STKs) in the catalytic loop, as well as the APE-2 residue (Trp in TKs, variable in STKs) in the activation loop. These residues correspond to positions 126, 128, and 165 in our MSA, respectively. Kinases which satisfy the conditions for TKs at all three positions were classified as TKs (10,345 raw sequences, 1,069 effective sequences) and those that satisfy the condition for STKs at position 126 and are non-overlapping with the TK condition at position 128 were classified as STKs (210,862 raw sequences, 22,893 effective sequences). The effective number of sequences in each class were calculated by summing over sequence weights, where each sequence was assigned a weight defined as the fraction of the number of sequences in the same class that are within 40% identity. In this way, we correct for the effects of phylogenetics in the calculation of sample size as well as other quantities (see below).

### Human kinome dataset

497 human kinase catalytic domain sequences were acquired from ref. 2 (excluding atypical kinases). These sequences were aligned using a Hidden Markov Model (HMM) that contains 259 columns (L = 259), which was constructed from the same MSA used to derive our Potts model. 447 human kinases remained after filtering-out sequences with 32 or more gaps.

### Classification of gatekeeper size

The designation of gatekeeper residues as “large” or “small” was based on sidechain van der Waals volumes^68^ (Fig. S6), where small gatekeepers have a volume of < 110 Å^3^ (Gly, Ala, Ser, Pro, Thr, Cys, Val) and large gatekeepers have a volume of > 110 Å^3^ (Asn, His, Ile, Leu, Met, Lys, Phe, Glu, Tyr, Gln, Trp, Arg).

### PDB dataset and conformational states

X-ray crystal structures of tyrosine, serine/threonine, and dual-specificity eukaryotic protein kinases in the PDB were collected from http://rcsb.org on July 30^th^, 2020. The protein sequences of 7,919 chains were extracted from 6,805 PDB files by parsing the SEQRES record and aligned to the MSA used to construct our Potts model, using a Hidden Markov Model (HMM). For this work, our classification of the active DFG-in and classical DFG-out conformational states is based on ref 24, which we describe in further detail in the Supplementary Information.

### Contact frequency differences

Each of the clustered PDB structures were converted to an adjacency matrix of binary contacts (1 for “in-contact”, 0 otherwise). A contact between residues *i* and *j* in structure *n* was assigned when their nearest heavy-atoms were detected within a distance *r*_*ij*_ (*n*) < 6 Å. The contact frequency 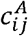 in cluster (e.g., active DFG-in) was calculated for each residue pair (*i,j*) by taking a weighted average over all instances of a contact in that cluster –

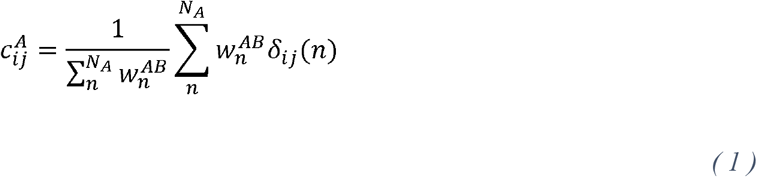

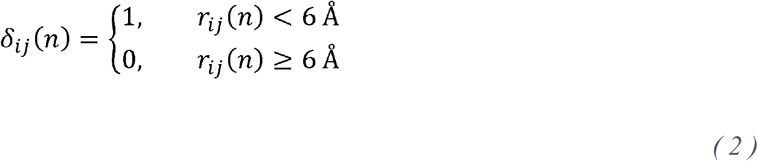

Where *N*_*A*_ is the number of PDB chains in cluster *A*, and weights were calculated with 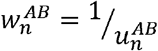 where 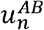 is the number of times the UniProt ID of structure *n* is found within either cluster *A* (i.e. Active) or cluster *B* (i.e. DFG-out). In this way, we have down-weighted contributions to the contact differences 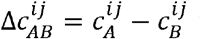 that are due to overrepresentation of specific kinases in the PDB clusters, with the goal of using contact differences to represent conserved features of the conformational transition across many different kinases. Alignment gaps and unresolved residues were accounted for by excluding these counts in the summations. Only |*i* − *j*| >4 were included in the calculation. The PDB clusters used to calculate these contact differences are described in the Supplementary Information.

### Potts model and threaded-energy calculation

Our Potts Hamiltonian was constructed from an MSA of protein kinase catalytic domains as previously described^67^. The Potts Hamiltonian *H*(*S*) takes the form –

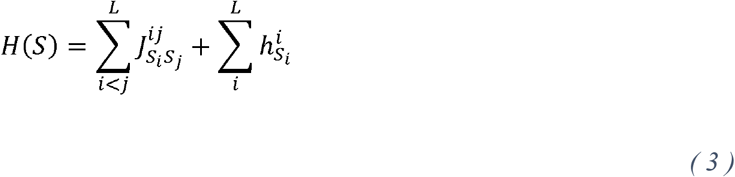

where *L* is the number of columns in the MSA (*L* = 259), *h* is a matrix of self-interactions or “fields”, and *j* is the coupling matrix which has the interpretation of co-evolutionary interactions between residues. The Potts threaded-energy penalty Δ*E*(*S*) for sequence *S* to undergo the conformational transition *A* ⟶ *B* is calculated using contact frequency differences between the two conformational ensembles^9^ –

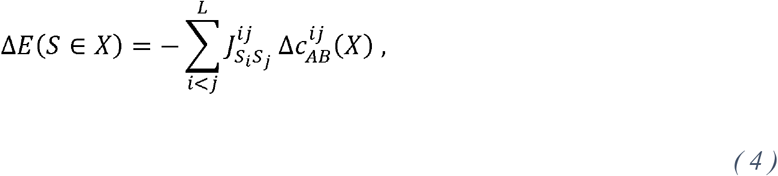

where *X* represents a class or family of sequences for which sequence *S* has membership, and 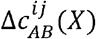 represents the contact frequency difference between conformations *A* and *B* observed only for other sequences belonging to class *X* (e.g. *X* ≡ *TKs*, or *X* ≡ *STKs*; upper and lower triangle of Fig. S2, respectively). As described previously^9^ the couplings 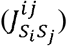 and fields 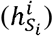 were transformed to the “zero-gauge” prior to calculating Δ*E*(*S*)

### Contributions to average shift in Δ*E*_*Potts*_ between STKs and TKs (ΔΔ*E*)

Where Δ*E*(*S*), is the Potts conformational penalty for sequence S to undergo the conformational change *A* ⟶*B*, we define ΔΔ*E* as the difference in average Δ*E* between two groups of sequences *X* and *Y*

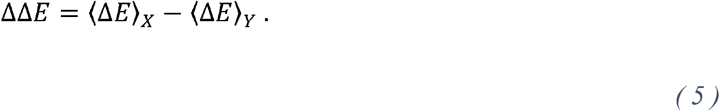

To help interpret ΔΔ*E* in a structural and coevolutionary context, we can write ΔΔ*E* as a sum over position pairs (*i,j*)

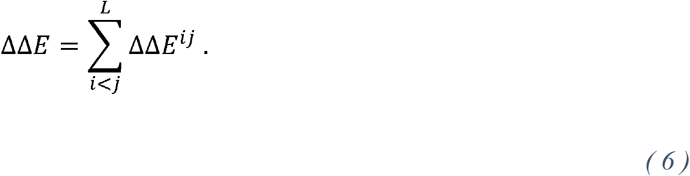

To evaluate this, we note that average Δ*E* can be expressed as a sum over position pairs

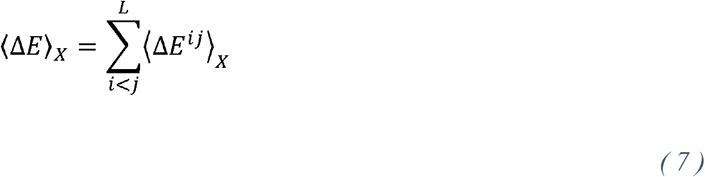

where ⟨Δ*E*^*ij*^⟩ is calculated as follows

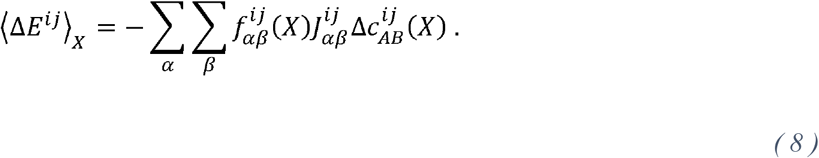

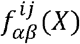 is the frequency (bivariate marginal) of residues *α* and *β* at positions *i* and *j* for sequences in the MSA which belong to group *X*, which we calculate after applying the MSA-derived phylogenetic weights described above. Finally, by substituting Eq. 7 back into Eq. 5, we show how ΔΔ*E* can be decomposed into contributions from individual residue pairs

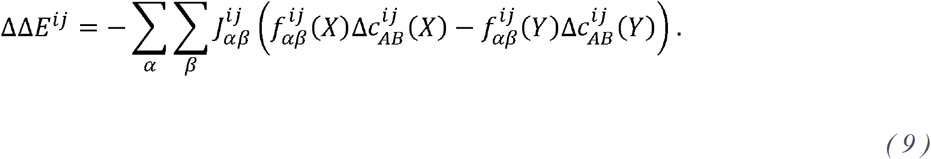

By viewing the largest (most positive) ΔΔ*E*^*ij*^ terms, where *X* ≡ *STKs*, and *Y* ≡ *TKs*, in Eq. 9, we are identifying position pairs that cause STKs to have higher penalties than TK in our Potts threading calculations for the active DFG-in to DFG-out conformational change (Fig. S5).

### Calculation of p-value for ΔΔ*E*

The quantity ΔΔ*E* is a difference between two averages, ⟨Δ*E*⟩_*STK*_ – ⟨Δ*E*⟩_*TK*_. Hypothesis testing to determine the statistical significance of this quantity was performed with respect to a null model where the populations of Δ*E* s for STKs and TKs, from which our samples were drawn, are indistinguishable. To this end, a p-value for was calculated for a t-statistic derived from Welch’s t-test^69^, where *s*, is the standard error of average Δ*E* –

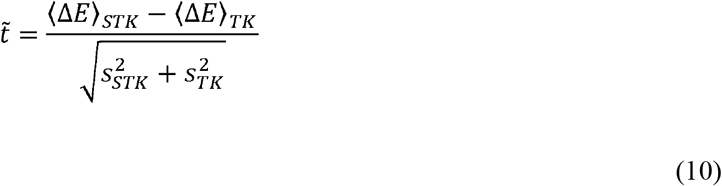

where the averages and standard errors are calculated after down-weighting each sequence as described above. This was done to lessen the effects of phylogenetic sampling bias in our MSA and ensure that ΔΔ*E* captures general differences between TKs and STKs, rather than specific TK or STK families.

The one-tailed p-value was calculated using the cumulative t-distribution function generated in python using the SciPy package^70^,

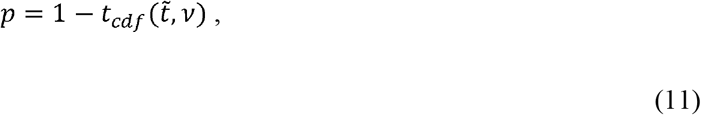

where the degrees of freedom for the t-distribution describing the combined population, *v*, was estimated via the Welch-Satterthwaite equation^69^ from the degrees of freedom of the two samples *v*_*STK*_ and *v*_*TK*_

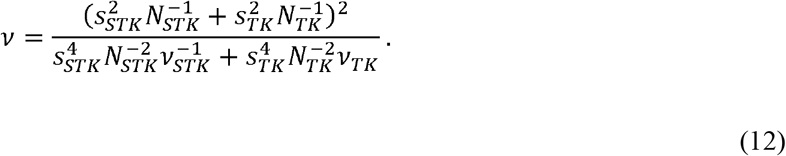

*N* represents the effective number of STKs or TKs, which is an unbiased count of sequences in each dataset that can be obtained by summing the sample weights (*N*_*STK*_ = 22,893, *N*_*TK*_ = 1069). From the calculation of ΔΔ*E* = ⟨Δ*E*⟩_*STK*_ − ⟨Δ*E*⟩_*TK*_ = 3.2, we determine the corresponding p-value to be less than 10^−15^, meaning it is highly unlikely for this large of a difference to be observed if the Δ*E*s for TKs and STKs were randomly drawn from the same distribution rather than distinct distributions.

### DDM Setup

The double decoupling method (DDM), also known as an “alchemical” method, was applied to compute absolute binding free energy 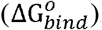 as shown in Eq. (13)^71,72^. This method computes the free energies of decoupling the inhibitor from the bulk solvent in the presence and absence of a receptor via a nonphysical thermodynamic cycle where the two end states are connected via the alchemical pathway. The starting holo-structures for absolute binding free energy (ABFE) calculations were taken from the available crystal structure. The absence of crystal structure prompted us to model the structure of the ligand into the active site of the kinase by superimposing over the binding pose of the available holo crystal structure.

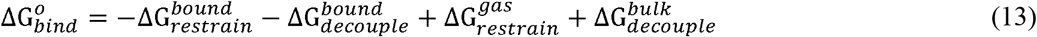

Decoupling of the ligand was achieved by first turning off the coulombic intermolecular interactions followed by Lennard-Jones intermolecular interactions from both the legs. This allows DDM to estimate the free energy, i.e., in the presence of protein 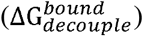 and absence of protein, i.e., in the bulk solvent, 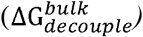 as shown in equations (14 and 15)

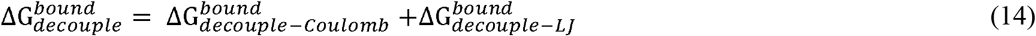

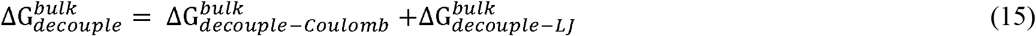

Substituting equations (14 and 15) into equation (16) yields the estimated 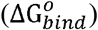 from DDM

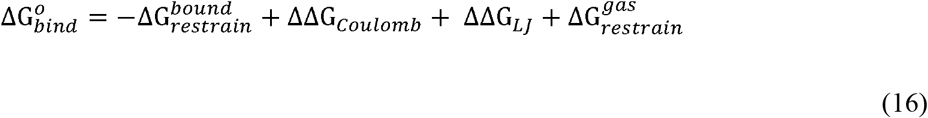

Where ΔΔG_*Coulomb*_ is the electrostatic energy contribution towards the total absolute binding free energy, and ΔΔG_*LJ*_ is the non-polar energy contribution.

In this study, depending on the system’s convergence, either 20 or 31 total *λ*s were used for decoupling the ligand from bulk solvent. For instance, either 5 *λ*s with Δ*λ* = 0.5 or 11 *λ*s with Δ*λ* = 0.1 were used for coulombic decoupling and 15 *λ*s with Δ*λ* = 0.1 or 20 *λ*s with Δ*λ* = 0.05 were used for decoupling Lennard-Jones interactions in the bulk solvent.

Similarly, depending on the convergence, either 30 or 42 total *λ*s were used for decoupling ligand bound to protein. For instance, 11 or 12 non-uniformly distributed *λ*s were used to restrain the ligand. Decoupling the coulombic interactions between ligand and protein was achieved by either using 4 *λ*s with Δ*λ* = 0.25 or 10 *λ*s with Δ*λ* = 0.1, whereas a large number of *λ*s were used for decoupling Lennard-Jones interactions, ie.,15 *λ*s with Δ*λ* = 0.1 or 20 *λ*s with Δ*λ* = 0.05 were used. The correction term developed by Rocklin and coworkers for treating charged ligands during DDM simulations were adopted^73^. In this regard, it is well documented that the use of a finite-sized periodic solvent box during DDM simulations can lead to non-negligible electrostatic energy contribution towards the calculated total ABFE. Thus, calculated 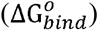 for charged ligand after addition of electrostatic correction term can be expressed as:

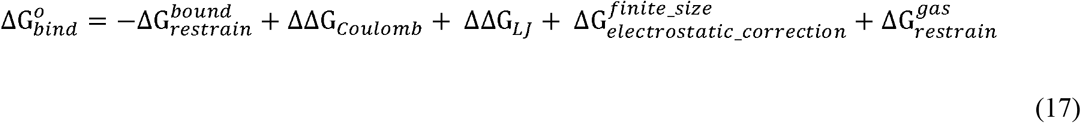

For a proper convergence during DDM simulations, the application of restrains is crucial. Herein, we have used six relative orthogonal restrains with harmonic potentials that include one distance, two angles, and three dihedral angles restrain between the ligand and the protein with a force constant of 10 kcal mol^−1^ Å^−2^ [deg^−2^]. At each *λ*, 10 to 30 ns of decoupling simulation via replica-exchange^74^ were obtained to compute the 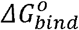 over a well-converged trajectory.

### MD Setup

In this study, molecular dynamics (MD) simulations were applied to compute the binding free energy simulations via DDM. GROMACS-2018.8^75^ was used as an MD engine for all simulations. The tleap module of AMBER16 was used to add the missing hydrogen atoms to the kinase enzymes. The system was solvated explicitly using TIP3P water boxes^76^ that extended at least 10 Å from the center of the system in each direction. The topology file for the kinase enzyme was created using the amber forcefield ff14SB^77^. The AM1-BCC charge model^78^ and general amber force field (GAFF2)^79^ were employed to parametrize different inhibitors used in this study. The overall charge of the system was maintained by adding a suitable number of counterions in each system. During the simulations, electrostatic interactions were computed using the particle mesh Ewald (PME) method^80^ with a cutoff and grid spacing of 10 and 1.0 Å, respectively. The NPT ensemble with a time step of 2 fs was used in the simulations..

## Data availability

Values of 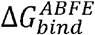 from all absolute binding free-energy simulations described in this work, including benchmarking calculations (94 simulations in total), are provided in the Supplementary Information. Potts Δ*E* s, type-II hit rates computed from ref. 7, the identity of gatekeeper residues and corresponding van der Waals volumes in Å^3^, and the classification of human kinases as TKs or STKs can be found in the Supplementary Table provided separately.

## Code availability

The Mi3-GPU^33^ source code required to reproduce the Potts model employed in this manuscript can be found at the following link: https://github.com/ahaldane/Mi3-GPU (v1.1).

## Acknowledgments

This research was supported by National Institutes of Health grant number R35-GM132090, and by NIH Computer Equipment Grant (OD020095). Gratitude is also expressed to the OWLSNEST high performance cluster at Temple University for its computing support in this project. We thank Shima Arasteh for helpful discussions related to kinase conformational states and alchemical free-energy simulations.

