## Supplementary Information for "Evolutionary Divergence in the Conformational Landscapes of Tyrosine vs Serine/Threonine Kinases"

### Supporting Information: "Evolutionary Divergence in the Conformational Landscapes of Tyrosine and Serine/Threonine Kinases"


Joan Gizzio[a,b1], Abhishek Thakur[a,b,1], Allan Haldane[a,c], Ronald M. Levy[a,b,2]

[a]Center for Biophysics and Computational Biology , Temple University, Philadelphia, Pennsylvania 19122

[b]Department of Chemistry, Temple University, Philadelphia, Pennsylvania 19122

[c]Department of Physics, Temple University, Philadelphia, Pennsylvania 19122

[1] J.G and A.T contributed equally to this work.

[2]To whom correspondence should be addressed.


#### I. Classifying protein kinase activation loop conformations and structural basis for the Potts threaded energy calculation

Before investigating the protein kinase conformational landscape using our sequence coevolutionary Potts model, it was necessary to first characterize the conformational states of interest using PDB x-ray crystal structures. We were interested specifically in the conformational change from the active state to the inactive, type-II-druggable DFG-out state usually associated with a large-scale reorganization of the activation loop.

Structural requirements for kinase catalytic activity are disrupted by rotation of the first three residues of the activation loop ("DF" of the conserved DFG motif and the variable residue preceding them defined as "X"). The conformational states of this small segment can be characterized by locations on the Ramachandran map (**B**eta-turn, right-handed **A**lpha-helix, **L**eft-handed alpha-helix) as well as the chi1 rotamer states of the DFG-Phe residue (gauche-**minus**, gauche-**plus**). In the PDB, there are eight clusters of XDF states identified by Dunbrack and co-workers (1) but only one of these (BLAminus) corresponds to the active conformation. The most populated inactive state in the PDB is BLBplus, which is often referred to as "Src-like inactive" in the literature. The classical DFG-out conformation, which is compatible with the binding of type-II inhibitors, corresponds to the BBAminus state.

In the active DFG-in (BLAminus) state, the ~20 residue long activation loop adopts an extended conformation which is maintained by contacts with the catalytic loop that "anchor" it at both ends, whereas the activation loop in the classical DFG-out state (BBAminus) tends to be observed in "folded" states with the N and C-terminal anchors compromised or broken. This phenomenon is detected from contact frequency differences between the active BLAminus (DFG-in) and inactive BBAminus (classical DFG-out) conformations (Figure S2). Incorporating these contact frequency differences into the Potts threaded energy calculation (see SI appendix) leads to a sequence and structure-based explanation for the generally high affinity of type-II inhibitors for TKs compared with STKs, wherein the free-energy required for the activation loop to reorganize from "extended" to "folded" is extremely important for controlling the conformational equilibrium between BLAminus and BBAminus.

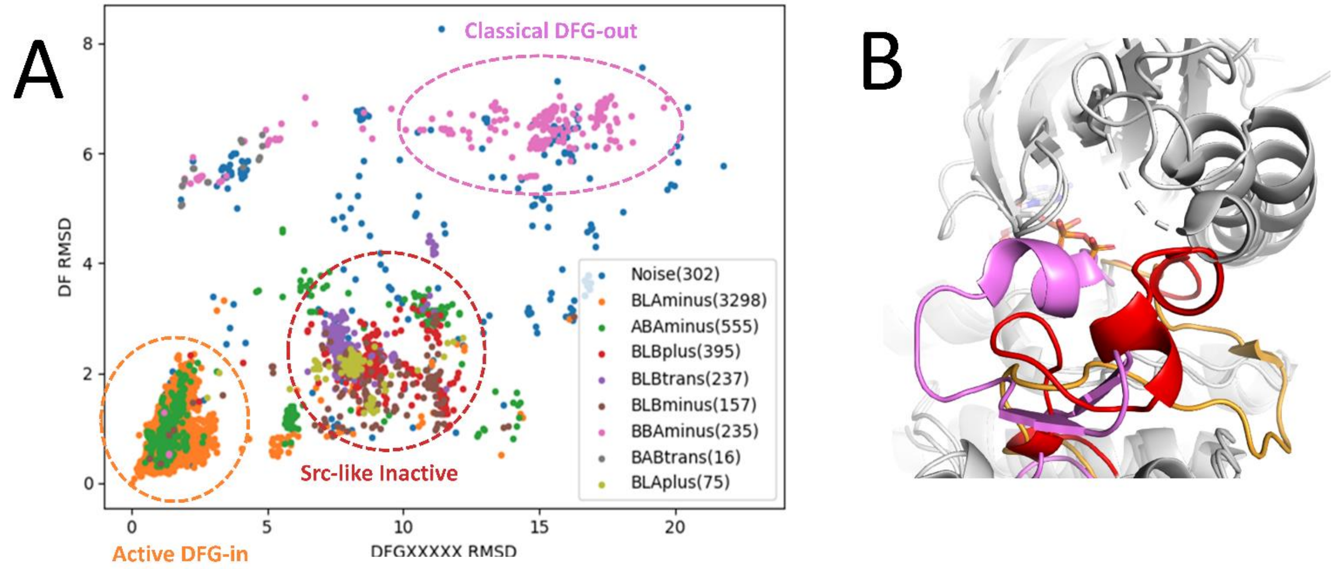


**Fig. S1.** Conformational landscape of the activation loop derived from the PDB. **(a)** Backbone torsions of the first three residues of the activation loop (XDF of the XDFG motif) broadly control folding of the activation loop into autoinhibitory conformations. The XDF dihedral classification (BLAminus, BLBplus, BBAminus, etc.), which informs the positioning of catalytic residues at the activation loop N-terminus, correlates with the spatial trajectory of the first five residues of the activation loop (DFGXXXXX) measured by RMSD with respect to catalytically primed Aurora A (PDB: 5DNR_A). Plotting the RMSD of the Asp and Phe residues of the DFG motif on the y-axis vs. RMSD of the DFGXXXXX residues on the x-axis captures three major conformational groups – an extended active-like conformation corresponding (circled orange) which is largely represented by the catalytically active BLAminus conformation, a partially folded DFG-in inactive conformation (circled red) which is mostly populated by BLBplus and commonly referred to as Src-like inactive, and the classical, folded DFG-out inactive conformation (circled pink) which is represented by the BBAminus state. **(b)** Cartoon representation of the three major conformational clusters created in PyMol.

**Active DFG-in (BLAminus).** The BLAminus state of the DFG motif (active DFG-in) is the only active conformation of the activation loop, in contrast to several other inactive DFG-in states (ABAminus, BLAplus, BLBplus, BLBminus, BLBtrans). In the active conformation, all structural requirements for catalytic activity are typically met: e.g. a complete hydrophobic spine, a salt bridge between $\beta$3-Lys $\rightarrow$ $\alpha$C-Glu, and an extended activation loop that ensures unobstructed substrate-binding, all of which have high correspondence with the BLAminus state.(1) Our analysis using software provided in ref. 1 identifies 3,643 structures in this conformation belonging to STKs and 625 structures belonging to TKs (4268 structures in total), which we used in the calculation of contact frequencies (Fig. S2).

**Classical DFG-out (BBAminus).** The DFG-out state is characterized by a "flip" of the conserved Phe to occupy the ATP binding pocket which is otherwise occupied by the conserved Asp in the active conformation. The classical DFG-out conformation is associated with the binding of type-II inhibitors which occupy the back pocket region opened up by the DFG-flip.(2) This conformation corresponds to the BBAminus rotamer state of these residues and it is the dominant DFG-out conformation associated with the binding mode of type-II inhibitors. This classical DFG-out or BBAminus conformation is correlated with a larger-scale conformational change of the activation loop that involves a ~18Å "folding" transition with respect to the active conformation. This conformational change is usually accompanied by the formation of secondary structure that obstructs the typical substrate binding surface (e.g. PDB ID: 2HIW, Fig. S1B). Our conformational analysis using ref. 1 identifies 224 structures in this conformational state belonging to STKs, and 286 structures belonging to TKs (510 structures in total). These structures were used to calculate contact frequencies which were then used to generate the contact differences plotted in Fig. S2 by subtracting them from contact frequencies in the active BLAminus state.

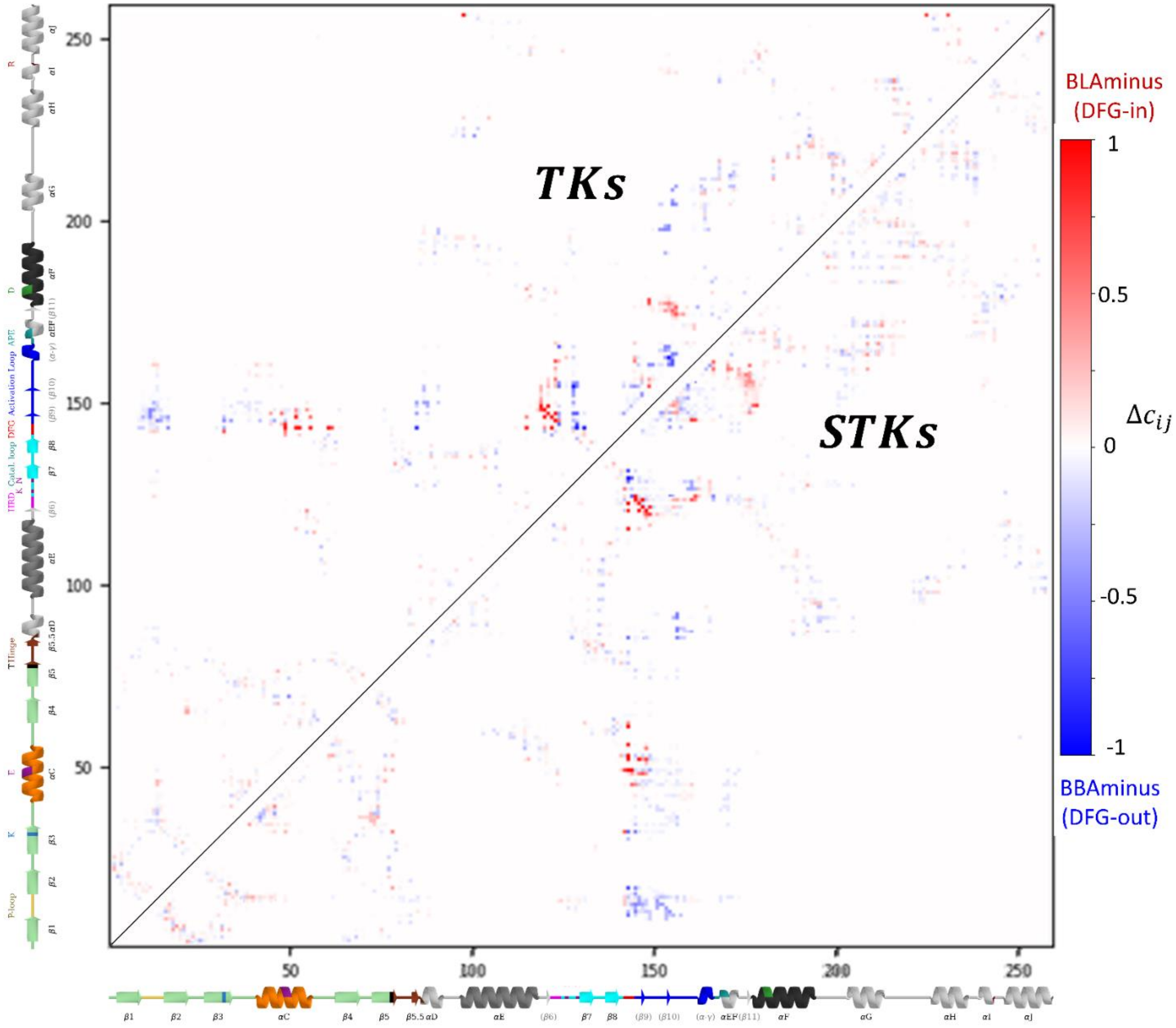


**Fig. S2.** Contact map depicting the contact frequency differences between the active DFG-in (BLAminus) and classical DFG-out (BBAminus) conformations. The contact frequency differences ($\Delta c_{ij} = c_{ij}^{BLA(-)} - c_{ij}^{BBA(-)}$) are explicitly involved in the calculation of $\Delta E_{Potts}$ (see *Methods* addendum). The upper triangle depicts the contact frequency difference matrix for kinases annotated in the PDB as ”Tyrosine Kinases”, meanwhile the lower triangle was calculated from kinases annotated as ”Serine/Threonine kinases”.

#### II. Benchmarking calculations for absolute binding free energy simulations

Accurate prediction of the absolute binding free energy difference between the Apo and Holo state of a protein is extremely important to achieve from force field-based molecular dynamics simulations.(3, 4) In this study, we used type-II inhibitors as probes to estimate $\Delta G_{reorg}$via absolute binding free energy (ABFE) simulations, which is the excess free-energy between experimentally determined binding affinity and $\Delta G_{bind}^{ABFE}$ calculated from ABFE (Eq. 3 in main text). Target kinases, i.e., MAPK14, CDK2 and JNK1 bound with type-I and $I^1/_2$ inhibitors in the active conformation states (where $\Delta G_{reorg}$ is expected to be close to zero) have been regularly used by the computational community as benchmark systems for absolute or relative binding free energy calculations.(5-10) Benchmarking calculations over multiple protein-ligand complexes show close agreement between calculated ($\Delta G_{bind}^{ABFE}$) and experimental ($\Delta G_{exp}^{o}$) terms (Table S2).

In the later part of benchmarking studies, we have included ABL1 bound to type-II inhibitors in the DFG-out fully folded state. As NMR studies have shown ABL1 has a $\Delta G_{reorg}$ of 1.2 kcal/mol(11), this system can be used to validate our ABFE simulation protocol. Our ABFE simulations with several type-II inhibitors bound to ABL1 in the DFG-out / activation loop folded conformation predicts an average $\Delta G_{reorg}$ of 1.26 kcal/mol (Table S7) which is consistent with the literature value. This result is an indicative that our estimates of $\Delta G_{reorg}$ are not significantly influenced by computational error (Table S1).

**Table S1:** Summary of benchmarking results described above, showing root mean square error (RMSE), mean unsigned error (MUE), and the average difference between experimental and computed binding free energy.

| Kinase | # Compounds | MUE | RMSE | Avg ($\Delta G_{exp}^{o} - \Delta G_{bind}^{ABFE}$) |
|---|---|---|---|---|
| CDK2 | 6 (type-I) | 0.71 | 1.03 | -1.89 |
| JNK1 | 5 (type-I) | 0.37 | 0.44 | -1.29 |
| MAPK14 | 9 (type-$1^1/_5$) | 0.47 | 0.64 | -0.2 |
| ABL1 | 6 (type-II) | 1.47 | 1.57 | 1.26 |

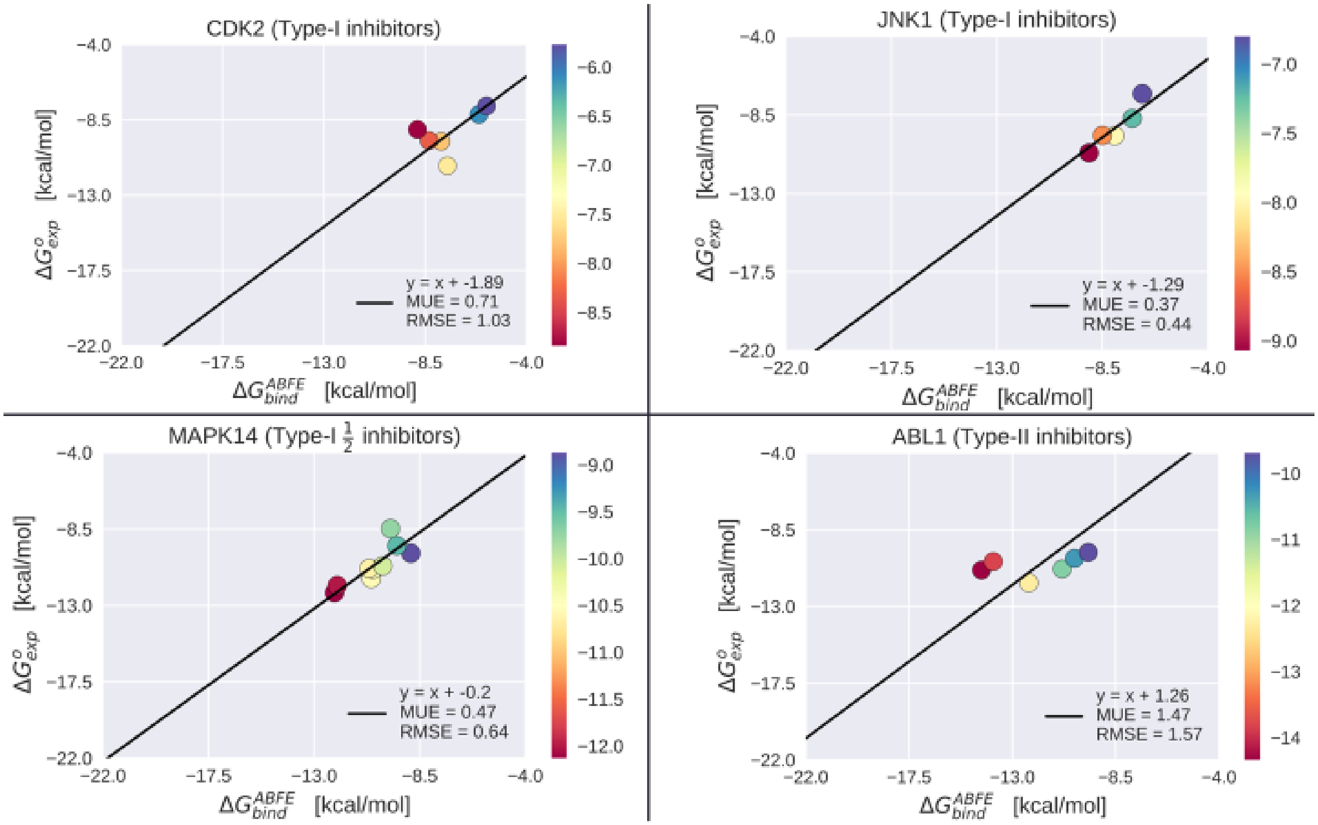


**Fig S3.** Benchmarking the absolute binding free energy results over 26 different co-crystallized inhibitors bound to CDK2, JNK1, MAPK14 in the active state and ABL1 in the inactive state.

#### III. Estimating the sequence-dependent $\Delta G_{reorg}$ over one inhibitor co-crystallized against multiple STKs and TKs via absolute binding free energy simulations

Over the decade Imatinib (Gleevec) has been co-crystallized against several other kinases besides ABL1 (e.g. DDR1, LCK, CSF1R, KIT, PDGFRA, and MAPK14) in the inactive conformation. Our ABFE calculations with Imatinib:MAPK14 complex, which is a STK, reveals a high reorganization penalty of 4-5 kcal/mol to adopt an inactive state, whereas none of the TKs co-crystallized against Imatinib show large $\Delta G_{reorg}$. We validated this result via an additional simulation with MAPK14, using instead a type-II inhibitor (BIRB-796) that is very potent for this kinase. Simulations with BIRB-796, which is a MAPK14 (p38a) selective inhibitor, show a very similar reorganization penalty to adopt the inactive conformation. This result confirms that the detected $\Delta G_{reorg}$ for MAPK14 fluctuates around the same value when using different type-II inhibitor probes with very different experimental potencies. Additional ABFE simulations of a few kinases co-crystallized against BIRB-796 also identify other STKs (MAPK9 and BRAF) as high penalty kinases, and PTK2B which is a TK as a low penalty kinase. Detailed results for these calculations are shown in Table S2.

**Table S2:** Calculated and experimental binding free energies for Imatinib and BIRB-796 against multiple kinases.

| Ligand | Kinase | Class# | $\Delta G^{o}_{exp}$ | $\Delta G^{ABFE}_{bind}$ |
|---|---|---|---|---|
| **Imatinib (STI)** | ABL1 (12) | TK | -10.80(13) | -10.82 ± 0.57 |
| | DDR1(14) | TK | -12.50(13) | -11.73± 1.22 |
| | LCK(15) | TK | -8.70(13) | -8.49 ± 1.28 |
| | CSF1R(16) | TK | -10.86(13) | -10.91 ± 1.29 |
| | KIT(17) | TK | -8.78(17) | -8.66 ± 1.43 |
| | PDGFRA | TK | -10.24(13) | -11.27 ± 0.27 |
| | MAPK14(18) | STK | -6.10(18) | -11.14 ± 0.75 |
| **BIRB-796 (B96)** | MAPK14(19) | STK | -12.75(13) | -17.00 ± 0.37 |
| | MAPK9(20) | STK | -11.10(13) | -19.88 ± 0.54 |
| | BRAF(21) | STK | -7.60(13) | -18.58 ± 0.35 |
| | PTK2B(22) | TK | -8.19(13) | -9.65 ± 0.13 |

### TK denotes Tyrosine kinase and STK denotes serine/threonine kinase, * Unit for $\Delta G_{exp}$ and $\Delta G_{calc}$ is kcal/mol.

#### IV. Identifying targets for absolute binding free energy simulations:

Our Potts threaded-energy calculations were used alongside experimental type-II binding data from the large-scale assay by Davis et al(13) to identify kinase targets that are likely to have very large or very small $\Delta G_{reorg}$. As described in the main text, all-atom molecular dynamics simulations to calculate absolute binding free-energies (ABFEs) of type-II inhibitors can be used alongside experimental binding affinities to calculate the free-energy cost for the kinase to reorganize to the DFG-out / folded activation loop conformation, via a simple relation (as described in Eq. 3 in the main text). This relation, $\Delta G_{reorg} = \Delta G^{o}_{exp} - \Delta G^{ABFE}_{bind}$, gives the free-energy cost to reorganize in physical energy units (kcal/mol) and can be used to approximate a scale for the Potts statistical energy differences provided one samples a sufficient range of $\Delta G_{reorg}$ and $\Delta E_{Potts}$. However, ABFE simulations are much more computationally demanding than the Potts threading calculation, which we sought to mitigate by choosing kinase simulation targets which are likely to provide a strong signal. We direct the reader to Table 1 in the main text, which contains the Potts penalties and type-II hit rates for the targets of interest. Fig. 1A and Fig. 5 in the main text provide the overall distributions for TKs and STKs, for comparison.

A significant challenge for our target selection was the limited availability of type-II inhibitors co-crystallized against STKs which have (a) Potts penalties and type-II hit rates that predict very high $\Delta G_{reorg}$, (b) experimental binding affinities available in the literature in the form of IC50, Ki, or Kd, (c) availability of protein-ligand co-crystallized structure/s and (d) for type-II inhibitor complex systems where the activation loop appears to have undergone a large-scale "folding" conformational change relative to the active "extended" conformation. STK complexes that satisfy all four criteria appear to be sparse, which seems consistent with the notion that kinases with large reorganization penalties are more difficult to crystalize in the DFG-out conformation(23). However, for some STKs with very high Potts threaded-energy penalties (e.g., MELK) there has been significant medicinal chemistry efforts to design potent type-II inhibitors and structurally characterize their complexes using x-ray crystallography. Using co-crystal structures that cover five different STKs with high Potts penalties (see main text, Table 1) and five different TKs with low Potts penalties, we were able to sample a wide range of $\Delta G_{reorg}$ from a total of 45 ABFE simulations covering 45 complexes and 10 different kinase targets. These simulations and subsequent calculations of $\Delta G_{reorg}$ for each kinase resulted in a wide range of values and strong correlation with the Potts statistical energy differences (see Fig. 5 in main text), allowing us to establish a scale for the Potts energies in kcal/mol. We have provided detailed results from the ABFE simulations of these 10 kinase targets in the following tables, and the corresponding plot of $\Delta G^{o}_{exp}$ vs $\Delta G^{ABFE}_{bind}$. In these plots, the average $\Delta G_{reorg}$ for each kinase (which averages out fluctuations in the data and more closely reflects the “real” value of $\Delta G_{reorg}$) is visualized as the y-intercept of a linear regression where the slope is constrained to one.

**Table S2:** Experimental and calculated binding free energies for type-I and II MELK inhibitors

| Kinase | Ligand | $\Delta G^{o}_{exp}$ | $\Delta G^{ABFE}_{bind}$ |
|---|---|---|---|
| MELK | 2 (24) | -6.30 | -10.49 ± 0.27 |
| | 3 (24) | -7.22 | -12.37 ± 0.3 |
| | 5* (24) | -8.38 | -13.47 ± 0.48 |
| | 4* (24) | -8.44 | -13.98 ± 0.22 |
| | 6 (24) | -9.71 | -17.14 ± 0.38 |
| | 7 (24) | -10.53 | -16.54 ± 0.52 |
| | 1* (25) | -5.18 | -8.06 ± 0.35 |
| | 5* (25) | -6.15 | -8.72 ± 0.13 |
| | 3* (25) | -7.34 | -9.71 ± 0.56 |
| | 9* (24) | -7.52 | -8.31 ± 0.32 |
| | 2* (25) | -7.75 | -8.45 ± 0.28 |
| | 6 (25) | -8.71 | -9.29 ± 0.51 |
| | 7* (25) | -10.14 | -10.86 ± 0.23 |

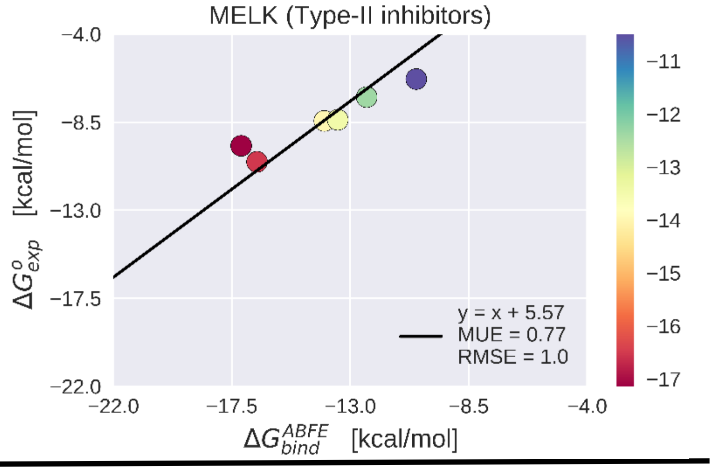


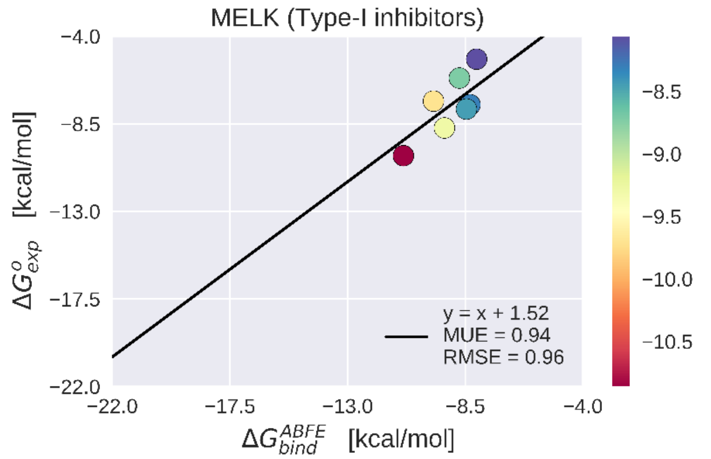


MELK (Maternal embryonic leucine zipper kinase) is STK that plays a very important role in the intracellular cell signaling pathways and multiple biological processes, i.e., apoptosis, cell cycle, tumorigenesis, etc. (26). TableS2 and the linear regression graph for MELK represent the experimental ($\Delta G^{o}_{exp}$) and calculated ($\Delta G^{ABFE}_{bind}$) for 7 and 6 type-I and II inhibitors, respectively, out of 23 data points in figure 3A. of main text.

**Table S3:** Experimental and calculated binding free energies for type-I and II IRAK4 inhibitors

| **Kinase** | **Ligand** | $\Delta G^{o}_{exp}$ | $\Delta G^{ABFE}_{bind}$ |
|---|---|---|---|
| **IRAK4** | Ponatinib (27) | -9.80 | -16.51 ± 0.33 |
| | HG-12-6(27) | -9.26 | -13.28 ±0.15 |
| | 8a* (28) | -11.88 | -13.32 ± 0.31 |
| | 16a* (28) | -11.64 | -11.80 ± 1.03 |
| | 15a* (28) | -10.77 | -10.04 ± 1.99 |
| | 1* (29) | -9.66 | -10.84 ± 0.94 |
| | 14 (29) | -8.08 | -9.85 ± 0.51 |
| | 12 (29) | -7.01 | -8.46 ± 0.99 |

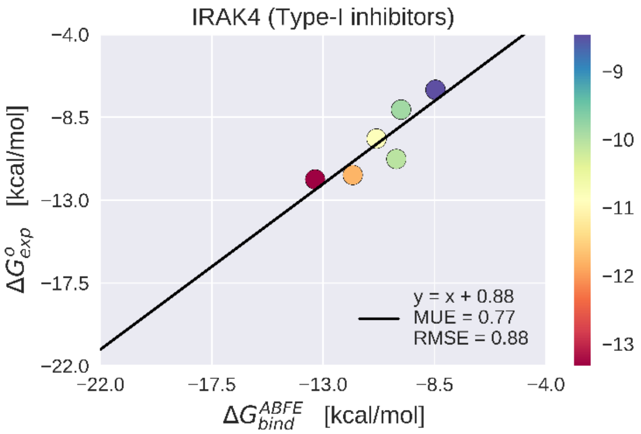


IRAK4 (Interleukin-1 receptor-associated kinase 4) is a STK that plays a major role in the immune response-related signaling pathways(30). A recent study has also indicated the involvement of IRAK4 in the immunopathogenesis against SARS-CoV-2 viral infection(31). TableS3 and the linear regression graph for IRAK4 represent the experimental ($\Delta G^{o}_{exp}$) and calculated ($\Delta G^{ABFE}_{bind}$) 6 and 2 type-I and II inhibitors, respectively, out of 23 data points in figure 3A of main text.

**Table S4:** Experimental and calculated binding free energies for type-I and II MAPK9 inhibitors

| Kinase | Ligand | $\Delta G^{o}_{exp}$ | $\Delta G^{ABFE}_{bind}$ |
|---|---|---|---|
| MAPK9 | HG-7-92-01 (32) | -6.83 | -11.96 ± 0.33 |
| | Sorafenib (33) | -7.01 | -15.26 ± 0.66 |
| | EXEL-2880 (34) | -8.43 | -13.84 ± 0.48 |
| | BIRB-76* (35) | -11.39 | -19.88 ± 0.54 |
| | 35* (36) | -9.40 | -10.06 ± 0.41 |

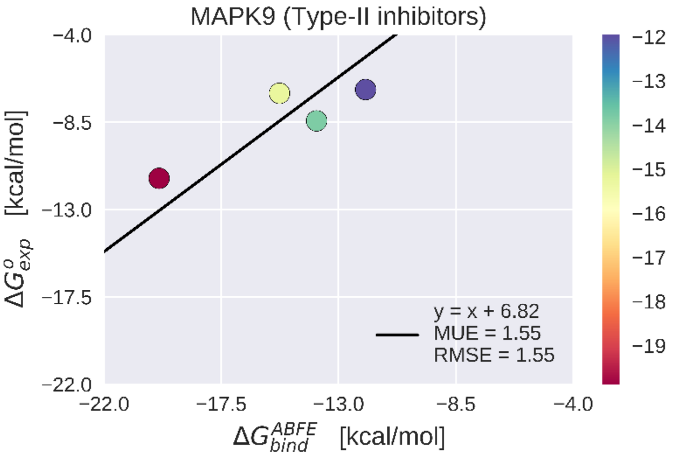


MAPK9 (Mitogen-activated protein kinase 9) also known as JNK2 (c-Jun N-terminal kinases) is a STK. It plays a vital role in multiple cellular processes such as expression of cytokines, stress responses, proliferation, and apoptosis(37). TableS4 and the linear regression graph for MAPK9 represent the experimental ($\Delta G^{o}_{exp}$) and calculated ($\Delta G^{ABFE}_{bind}$) for 1 and 4 type-I and II inhibitors, respectively, out of 23 data points in figure 3A of main text.

**Table S5:** Experimental and calculated binding free energies for type-I and II CDK2 inhibitors

| Kinase | Ligand | $\Delta G^{o}_{exp}$ | $\Delta G^{ABFE}_{bind}$ |
|---|---|---|---|
| CDK2 | K03859 (36) | -6.83 | -15.03 ± 0.27 |
| | Sorafenib (13) | -6.91 | -15.1 ± 0.25 |
| | K03861* (36) | -9.97 | -16.27 ± 0.51 |
| | 1H1S (38) | -11.25 | -7.51 ± 0.36 |
| | 1OIY (39) | -9.79 | -7.79 ± 0.19 |
| | 1OI9(39) | -9.74 | -8.33 ± 0.15 |
| | 1OIU(39) | -9.08 | -8.84 ± 0.14 |
| | 1H1Q(38) | -8.18 | -6.11 ± 0.15 |
| | 1H1R(38) | -7.67 | -5.77 ± 0.18 |

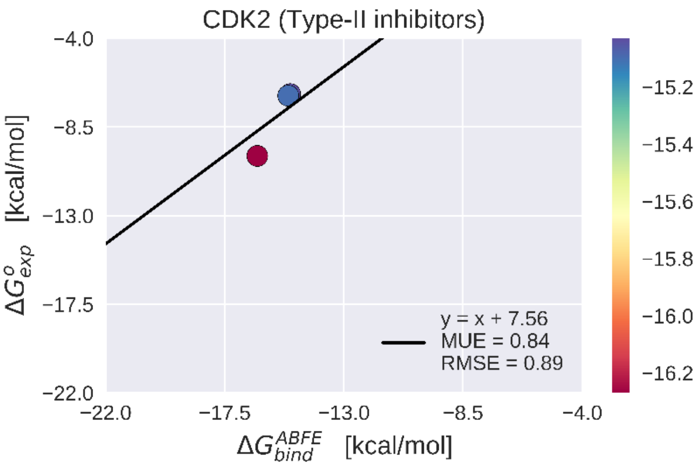


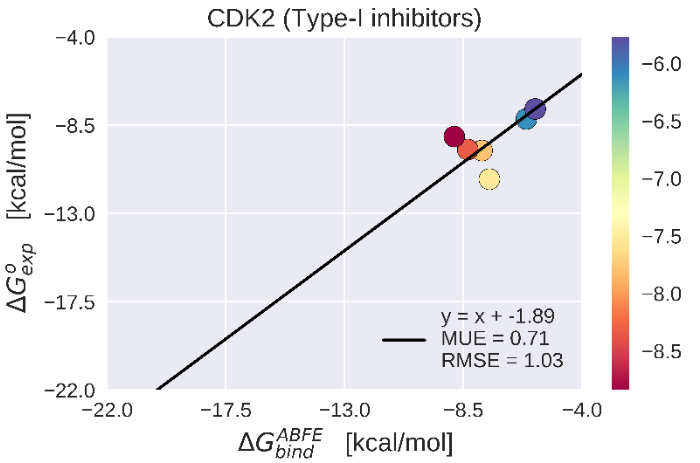


CDK2 (Cyclin-dependent kinase 2) is also known as cell division protein kinase 2. It is a STK that plays a major role in cell cycle regulations of G1/S and S/G2 phase transitions (40). TableS5 and the linear regression graph for CDK2 represent the experimental ($\Delta G^{o}_{exp}$) and calculated ($\Delta G^{ABFE}_{bind}$) 6 and 3 type-I and II inhibitors, respectively, out of 23 data points in figure 3A of the main text.

**Table S6:** Experimental and calculated binding free energies for type-I and II BRAF inhibitors

| Kinase | Ligand | $\Delta G^{o}_{exp}$ | $\Delta G^{ABFE}_{bind}$ |
|---|---|---|---|
| BRAF | 4*(41) | -11.79 | -19.45 ± 0.31 |
| | 8b*(42) | -11.03 | -15.4 ± 0.24 |
| | 6d*(43) | -10.82 | -16.72 ± 0.62 |
| | BAY89006*[7] | -10.46 | -14.72 ± 0.24 |
| | 5*(44) | -9.78 | -13.60 ± 0.35 |
| | RAF-265*(45) | -9.77 | -15.25 ± 0.48 |
| | Ponatinib*(46) | -8.95 | -18.03 ± 0.23 |
| | BIRB-76*(21) | -7.6 | -18.58 ± 0.35 |
| | 18*(47) | -14.37 | -12.73 ± 0.05 |
| | 20(47) | -13.19 | -13.78 ± 0.24 |
| | 23(47) | -13.16 | -14.83 ± 0.03 |

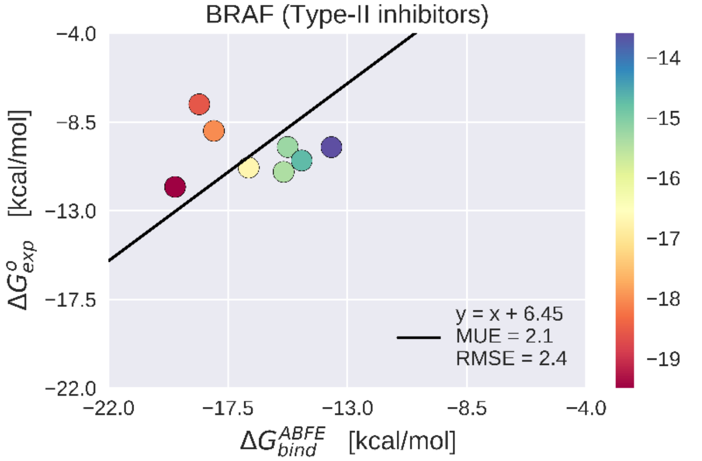


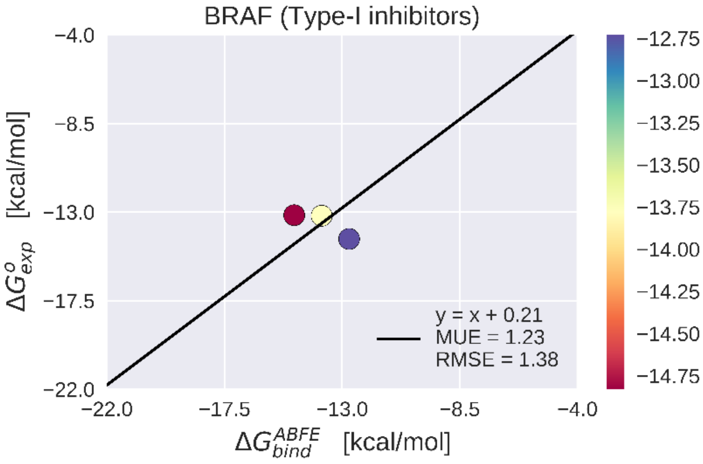


BRAF is a STK that plays a very important role in mediating signals that are responsible for promoting cell growth and proliferation (48). TableS6 and the linear regression graph for BRAF represent the experimental ($\Delta G^{o}_{exp}$) and calculated ($\Delta G^{ABFE}_{bind}$) for 3 and 8 type-I and II inhibitors, respectively, out of 23 data points in figure 3A of the main text.

**Table S7:** Experimental and calculated binding free energies for type-II ABL1 inhibitors

| Kinase | Ligand | $\Delta G^{o}_{exp}$ | $\Delta G^{ABFE}_{bind}$ |
|---|---|---|---|
| ABL1 | 4(49) | -11.62 | -12.28 ± 1.19 |
| | 406(8)*(49) | -10.87 | -14.34 ± 0.48 |
| | Imatinib*(12) | -10.80 | -10.83 ± 0.55 |
| | 44*(50) | -10.37 | -13.83 ± 0.31 |
| | 2(49) | -10.18 | -10.29 ± 1.31 |
| | 35(50) | -9.83 | -9.69 ± 0.32 |

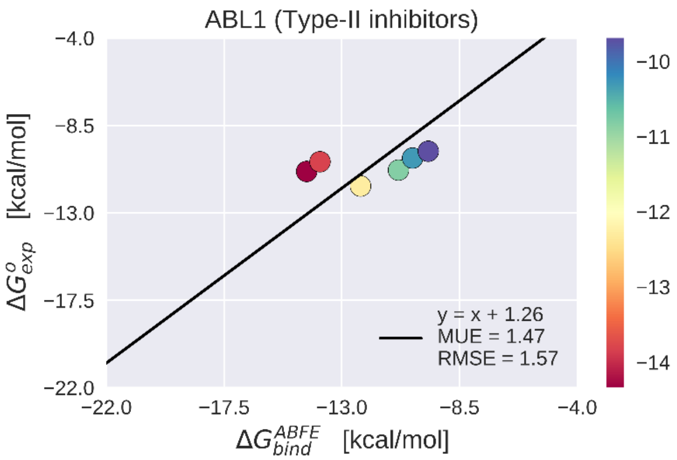


ABL1 is a TK protein and is mainly responsible for chronic myelogenous leukemia (CML)(51) TableS7 and the linear regression graph for ABL1 represent the experimental ($\Delta G^{o}_{exp}$) and calculated ($\Delta G^{ABFE}_{bind}$) of 6 type-II inhibitors out of 20 data points in figure 3B

**Table S8:** Experimental and calculated binding free energies for type-II DDR1 inhibitors

| Kinase | Ligand | $\Delta G^{o}_{exp}$ | $\Delta G^{ABFE}_{bind}$ |
|---|---|---|---|
| **DDR1** | Imatinib*(14) | -11.91 | -11.73 ± 1.22 |
| | 7c*(52) | -11.27 | -9.88 ± 0.28 |
| | 7m(52) | -9.78 | -8.25 ± 0.22 |
| | 6c*(53) | -8.89 | -10.98 ± 1.07 |
| | 6h(53) | -8.85 | -11.25 ± 1.56 |

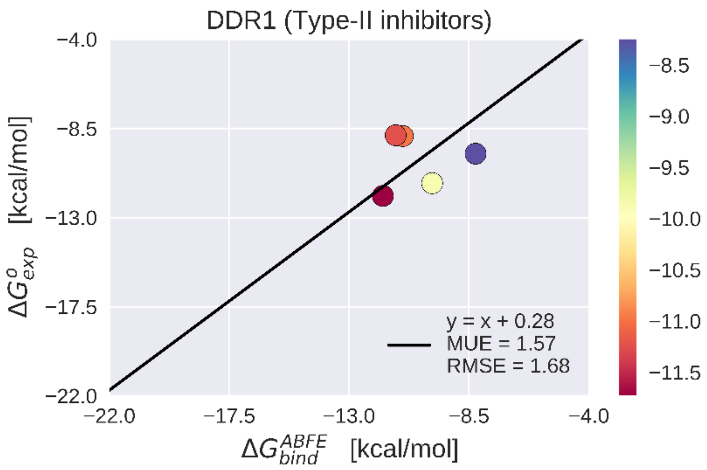


DDR1 (Discoidin domain receptor, family member-1) also known as CD167a (cluster of differentiation 167a) is a TK protein and is mainly responsible for cell regulation, growth, and metabolism (54). TableS8 and the linear regression graph for DDR1 represent the experimental ($\Delta G^{o}_{exp}$) and calculated ($\Delta G^{ABFE}_{bind}$) for 5 type-II inhibitors out of 20 data points in figure 3B.

**Table S9:** Experimental and calculated binding free energies for type-II LCK inhibitors

| Kinase | Ligand | $\Delta G^{o}_{exp}$ | $\Delta G^{ABFE}_{bind}$ |
|---|---|---|---|
| LCK | 10b*(55) | -12.70 | -11.95 ± 0.78 |
| | 19*(55) | -12.42 | -13.33 ± 0.92 |
| | 16(55) | -11.33 | -13.9 ± 0.36 |
| | Imatinib*(15) | -8.87 | -10.18 ± 0.56 |

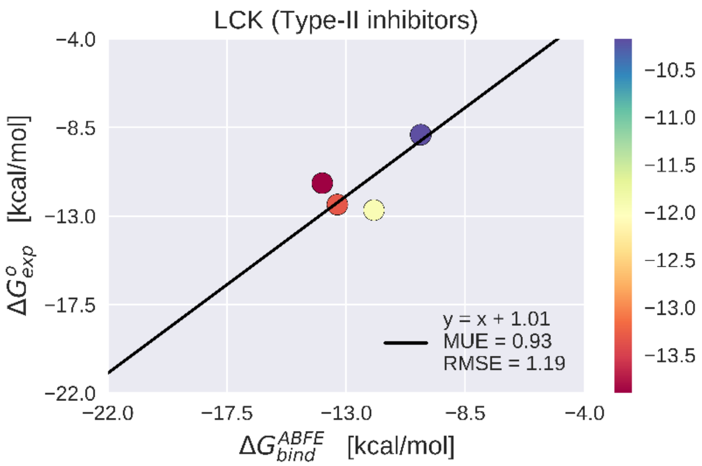


LCK (Lymphocyte-specific protein) is a TK protein that belongs to the Src family and is responsible for the intracellular signaling pathways. (56) TableS9 and the linear regression graph for LCK represent the experimental ($\Delta G^{o}_{exp}$) and calculated ($\Delta G^{ABFE}_{bind}$) 4 type-II inhibitors out of 20 data points in figure 3B of the main text.

**Table S10:** Experimental and calculated binding free energies for type-II TIE2 inhibitors .

| Kinase | Ligand | $\Delta G^{o}_{exp}$ | $\Delta G^{ABFE}_{bind}$ |
|---|---|---|---|
| TIE2 | 1b*(57) | -12.29 | -11.83 ± 0.44 |
| | 15*(58) | -10.92 | -10.68 ± 0.34 |
| | 2*(58) | -10.61 | -10.69 ± 0.45 |
| | 42(58) | -9.31 | -10.24 ± 0.05 |
| | 46(58) | -8.74 | -7.12 ± 0.14 |

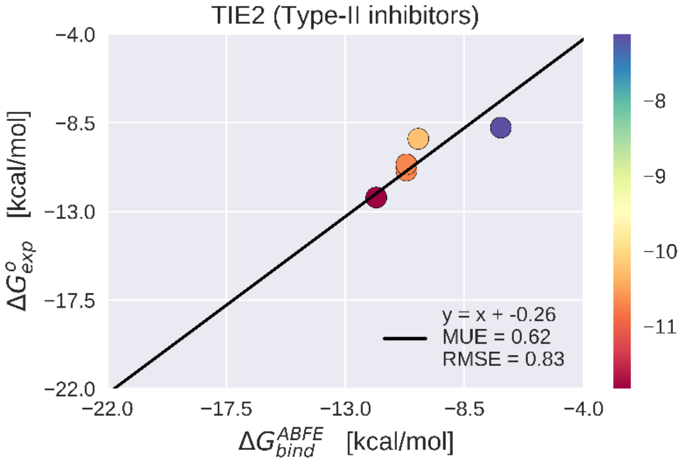


TIE2 is an angiopoietin receptor that is classified as a TK protein. This protein is responsible for various signaling pathways in the endothelial cells. (59) TableS10 and the linear regression graph for TIE2 represent the experimental ($\Delta G^{o}_{exp}$) and calculated ($\Delta G^{ABFE}_{bind}$) of 5 type-II inhibitors out of 20 data points in figure 3B of main text.

**Table S11:** Experimental and calculated binding free energies for type-II NTRK2 inhibitors

| Kinase | Ligand | $\Delta G^{o}_{exp}$ | $\Delta G^{ABFE}_{bind}$ |
|---|---|---|---|
| **NTRK2** | EX429*(60) | -10.82 | -11.85 |
| | GW2580*(60) | -10.16 | -12.57 |

NTRK2 (Neurotrophic Tyrosine Kinase, Receptor, Type 2) also known as TrkB (Tropomyosin receptor kinase B) or BDNF/NT-3 growth factors receptor is a TK protein. It is a well know receptor for its neuroprotective functions and its dysregulation has been associated with multiple neurological and neurodegenerative conditions (61). TableS11 and the linear regression graph for NTRK2 represent the experimental ($\Delta G^{o}_{exp}$) and calculated ($\Delta G^{ABFE}_{bind}$) for 2 type-II inhibitors out of 20 data points in figure 3B of main text.

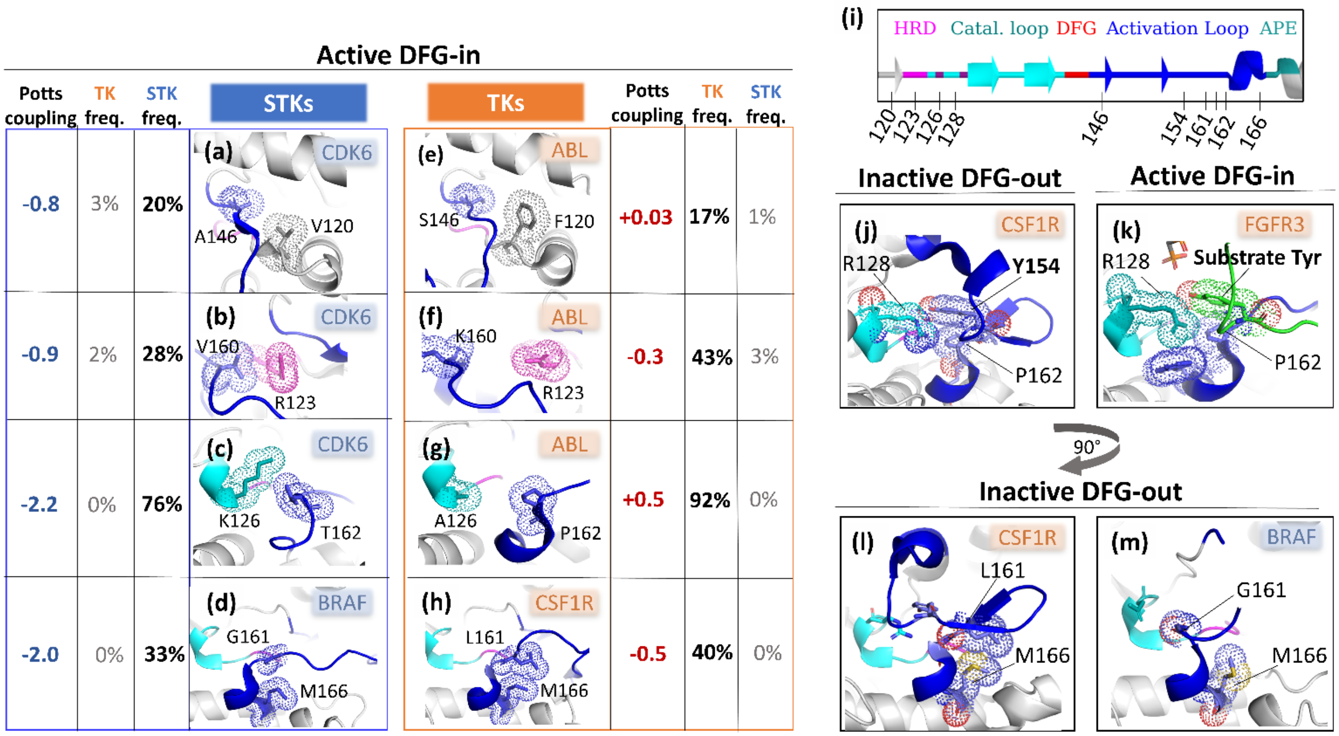


**Fig. S4. Analysis of the top 5 residue interactions.** $\Delta\Delta E$ (see Figure 5 in the main text) was decomposed to identify the strongest contributing pairs of positions (see *Methods*), with the idea that these pairs form interactions in the 3D structure that are important for controlling the conformational equilibrium. The residue pairs shown in this figure belong to the top 5 $\Delta\Delta E^{ij}$ (except for (128, 154), which is the $19^{\text{th}}$ top contributing pair). For each position pair (a-h), the residue identities which make the largest contributions are shown. Location of each position along the kinase primary structure with consistent coloring is shown in (i) for reference. (j-k) shows the substrate mimicking behavior of the TK DFG-out activation loop, with the trans-autophosphorylating FGFR3 structure for reference (substrate activation loop colored green). (l) The interaction between positions (161, 166) forms part of the activation loop C-terminal bulge in TKs which, according to the Potts model, decreases the energetic cost for DFG-out relative to STKs (m) where formation of this interaction pair is constrained to the active conformation and transition to DFG-out is penalized by breakage of this contact. PDBs: 1XO2 chain B (active CDK6), 2G2I chain A (active ABL), 4MNE chain B (active BRAF), 5HID chain A (inactive BRAF), 3LCD chain A (active CSF1R), 4R7I chain A (inactive CSF1R, 6PNX (trans-autophosphorylating FGFR3 dimer).

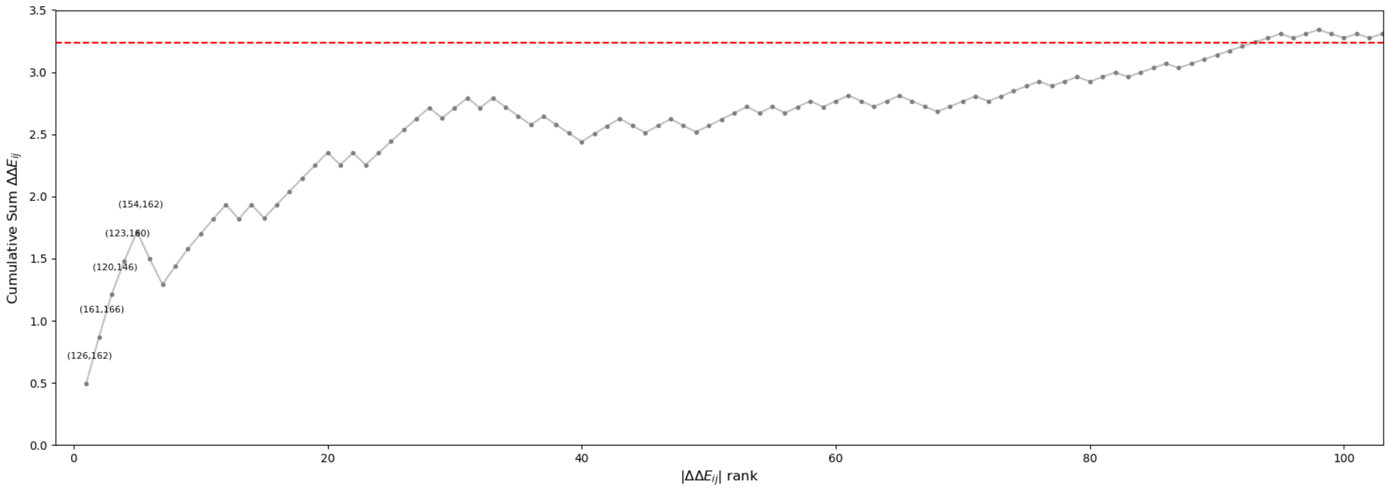


**Fig. S5. Cumulative contributions to ΔΔ$E$ from individual position pairs.** The position pair contributions to ΔΔ$E$ (see *Methods* in the main text) were ordered from largest to smallest magnitude. The top pairs which are discussed in the main text and illustrated in Figure S4 are annotated.

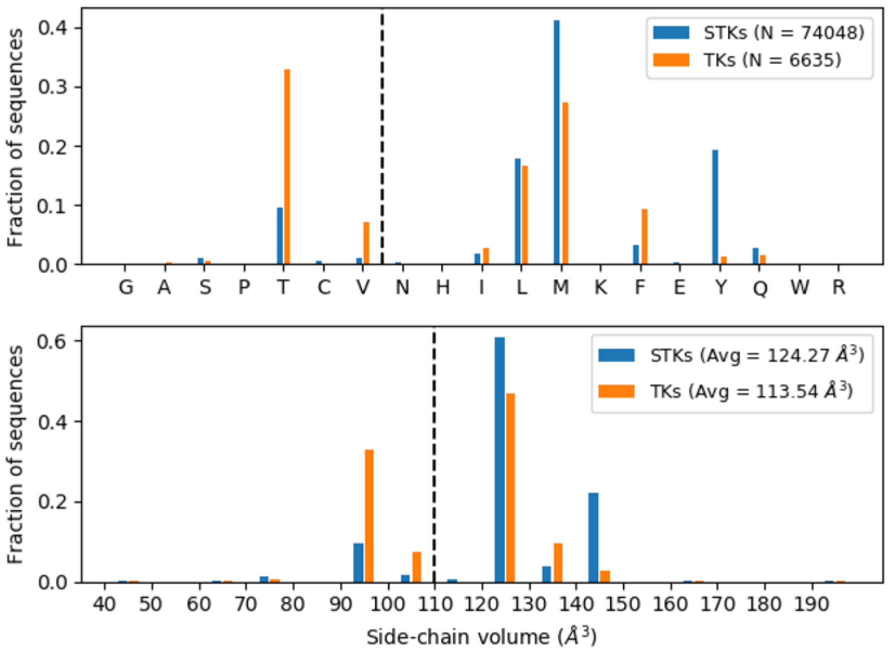


**Fig. S6.** Thousands of STK and TK sequences (see legend) were analyzed to determine the identities of the gatekeeper residues. Distribution of gatekeeper residues (top) with bins ordered (left to right) according to increasing sidechain volume(64) shows that STKs tend to have Met as a gatekeeper, whereas TKs tend to have Thr. Alternatively, one can look at the distribution of gatekeepers in terms of sidechain volume (bottom) which shows TKs have a greater tendency towards small gatekeepers in comparison with STKs.
